# The impact of the spatial heterogeneity of resistant cells and fibroblasts on treatment response

**DOI:** 10.1101/2021.06.01.446525

**Authors:** Masud M A, Jae-Young Kim, Cheol-Ho Pan, Eunjung Kim

## Abstract

A long-standing practice in the treatment of cancer is that of hitting hard with the maximum tolerated dose to eradicate tumors. This continuous therapy, however, selects for resistant cells, leading to the failure of the treatment. A different type of treatment strategy, adaptive therapy, has recently been shown to have a degree of success in both preclinical xenograft experiments and clinical trials. Adaptive therapy is used to maintain a tumor’s volume by exploiting the competition between drug-sensitive and drug-resistant cells with minimum effective drug doses or timed drug holidays. To further understand the role of competition in the outcomes of adaptive therapy, we developed a 2D on-lattice agent-based model. Our simulations show that the superiority of the adaptive strategy over continuous therapy depends on the local competition shaped by the spatial distribution of resistant cells. Cancer cell migration and increased carrying capacity accelerate the progression of the tumor under both types of treatments by reducing the spatial competition. Intratumor competition can also be affected by fibroblasts, which produce microenvironmental factors that promote cancer cell growth. Our simulations show that the spatial architecture of fibroblasts modulates the benefits of adaptive therapy. Finally, as a proof of concept, we simulated the outcomes of adaptive therapy in multiple metastatic sites composed of different spatial distributions of fibroblasts and drug-resistant cell populations.

## 1. Introduction

The current standard of care for the treatment of cancer patients is based on continuous therapy (CT) using the maximum tolerated dose of cancer drugs with the aim of eradicating tumors by killing the maximum number of drug-sensitive cancer cells. Despite the impressive initial tumor responses under CT, drug resistance inevitably develops in advanced metastatic solid cancers because CT often selects for drug-resistant cell populations [1,2]. For example, a majority of patients with metastatic melanomas treated continuously with BRAF-MEK inhibitor experienced a progression of the disease over 11–15 months [3,4]. The development of resistance is known to be a combined consequence of the responses from factors that include intratumor heterogeneity [5,6], limited drug penetration due to physical barriers [7], and the tumor microenvironment [8–10]. Thus, the exploitation of the intratumor competition between heterogeneous cancer cells and the modulation of the tumor microenvironment to bias the selective pressure towards the sensitive cells have the potential to delay the emergence of resistance.

From an ecological and evolutionary perspective, the net growth rate of a population composed of multiple species is determined by the intrinsic growth rate, death rate, and density-dependent limitations—when multiple species compete for the same resources in a closed environment [11]. This ecological principle implies that the net growth of a tumor cell population can be modulated by inhibiting the intrinsic growth rate of drug-sensitive cells, by increasing sensitive cell deaths, and by modulating the density-dependent growth limitations of drug-resistant cell populations. Because drug resistance often comes with a fitness cost [12,13], treatment breaks may provide sensitive cells with a higher net growth rate than the resistant cell population. When the intrinsic growth rates of both cell populations are same, the only way to modulate the growth of resistant cells is to increase density-dependent limitations. Adaptive therapy (AT) is based on this ecological principle of competition between tumor cells to limit their growth [14]. If kept in a tolerable range, tumor burden is not always lethal [14,15]. Thus, the objective of AT is to maintain a tolerable tumor burden as long as possible by using treatment holidays and reduced dosing [14]. For example, under AT, a patient is treated with therapy from the diagnosis until the tumor burden falls to a fraction of the initial cell population (e.g., 50% of the initial burden [16]). The goal is to reduce the cell population to an acceptable level that has sufficient sensitive cells to maintain density-dependent competitive stress on the growth of resistant cells. Then, a treatment break is scheduled to allow the remaining sensitive cells to grow and to limit the growth of the resistant cell population by leveraging competition. Once the total cell population is back to the initial level, a treatment is administered again. This on–off treatment cycle is repeated until the tumor progresses. This adaptive therapy strategy has been shown to have some degree of success in both preclinical experiments [17,18] and a clinical trial [16]. In particular, a clinical trial for prostate cancer therapy showed that adaptive therapy can delay disease progression for 27 months by using only a 53% cumulative drug rate compared to CT [16].

Several mathematical and computational models have been developed to compare AT with CT in various scenarios. Two key terms in this regard are time to relapse and tumor progression (TTP), which is the time at which the tumor volume exceeds 120% of the the initial volume; time gain (TG) which is defined as: TG = TTP in AT – TTP in CT. Gallaher et al. developed an off-lattice agent-based model to simulate the impact of heterogeneity and space on AT outcomes. They reported an extension of TTP of about one year under AT compared to CT (CT: 400 days vs. 700 days) [19]. Gatenby et al. developed a model consisting of five types of cells with differential drug responses and showed that tumor cells under CT grow to a carrying capacity by about 2400 days, while under AT, the tumor burden was kept under control at 20% of the carrying capacity [14]. A mathematical model in [16] showed that the on–off cycling rate of treatments depends on cell–cell competition and initial tumor cell population composition [16], where the threshold for treatment breaks was 50% of the initial tumor burden. A different threshold for treatment breaks was considered by Hansen and Read [20]. This study further demonstrated that a 20% reduction threshold resulted in more delayed progression than a 50% reduction for different degrees of initial resistance [20]. Some studies identified critical factors that determined the TG of AT. In the case of melanoma, the initial tumor burden, growth rate, switching rate, and competition coefficient were identified as crucial parameters for deciding the TG of AT by using CT [21]. The initial proportion of resistance is another contributing factor. Strobel et al. showed that a 1% initial resistance delayed the progression by up to 211 days for an initial burden of 75%, while 10% resulted in almost no TG [22]. A game-theoretical model was used to propose a combination of strategies for AT [23,24]. Recently, Viossat and Roble [25] provided theoretical conditions for the maximization of the benefits of AT. In particular, they provided an explicit formula for TG under AT, which included the intensity of competition between drug-sensitive and drug-resistant cells, the most critical factor.

Because the competitive stress experienced by each cell depends on its neighborhood in solid tumors, spatial models would be more suitable for exploring the consequences of spatial heterogeneity and treatment for tumor growth [26]. Agent-based models have shown that even lower doses can limit tumor growth if resistant cells are spatially restricted by sensitive cells [19,27]. In tumors with cells of a varied range of sensitivity, AT resulted in trapping of the resistant cells by the sensitive cells, thus limiting the growth of the tumor [19]. Tumors with spread randomly resistant cells were reported to grow much faster than tumors with resistant cells that were clustered together [27,28]. AT has even been found to delay progression in the absence of fitness costs [27,28], which, however, are assumed to be a key element for the success of AT [12,13].

Furthermore, the tumor microenvironment can be used to modulate tumor growth and competition. For example, fibroblasts are known to act as a local moderator of individual cancer cells’ growth and migration by producing growth factors and an extracellular matrix [29–34]. The spatial heterogeneity in tumor growth could be affected by fibroblast locations in the tumor [31]. It was demonstrated that physical proximity to fibroblasts determines tumor cell survival under therapy. Tumor cells that are close to fibroblasts can survive longer under therapy due to the fibroblast-mediated elevation of the threshold of drug concentration required for cell death and the lower rate of drug activity due to the physical barrier against drug penetration (i.e., collagen) generated by fibroblasts.

Our study aims to investigate the impacts of the spatial distributions of both resistant cells and fibroblasts on therapeutic outcomes. We developed a 2D on-lattice agent-based model (ABM) and inspected tumor growth subject to three different initial resistant cell configurations—namely, clumped, random, and uniform—to explore the impact of the spatial arrangement of cells. We simulated both AT and CT and compared the TTP. Furthermore, we explored the impact of the carrying capacity and cell migration rate on cell–cell competition and treatment outcomes. For the fibroblast locations, we considered two different proximities to resistant cells: overlapping with resistant cells and close to resistant cells. Finally, we simulated tumor growth in a virtual patient with four metastatic sites composed of different migration rates and different spatial distributions of fibroblasts and resistant cell populations. In these simulations, we assumed that one of the metastatic sites was not detected at the beginning of the treatment. The tumor progression was determined by utilizing two criteria: the sum of tumor burden and the emergence of a new metastatic lesion driven by the growth of an initially undetected metastatic lesion.

## 2. Materials and Methods

To study how a resistant cell population modulates treatment response, we considered a 2D on-lattice agent-based model of a small primary tumor or a metastatic lesion. For simplicity, we assume that a tumor cell population can be classified into two types of cells: drug-sensitive (S-cell) and drug-resistant (R-cell). We denote the total cell population, S-cell population, and R-cell population at time *t* with *N*(*t*), *S*(*t*), and *R*(*t*), where *N*(*t*) = *S*(*t*) + *R*(*t*).

### 2.1. Initial and Boundary Conditions

We assume that a percentage *f*_*R*_ of the initial cells (*N*(0)) are resistant (i.e., 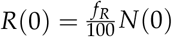 and 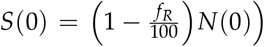. Initially, a total of *S*(0) cells are randomly dispersed over the domain, while a total of *R*(0) cells were placed in three different dispersion patterns—random, uniform, or clumped—in the domain.[35]. In the clumped case, all of the R-cells were randomly dispersed in a square centered in the middle of the domain, where the same number of R-cells were randomly dispersed over the whole domain in the random case. On the other hand, in the uniform case, all of the R-cells were manually placed to maximize the distance between R-cells over the whole domain. Please refer to the Figure 1 for the three types of cell configurations.

**Figure 1.**
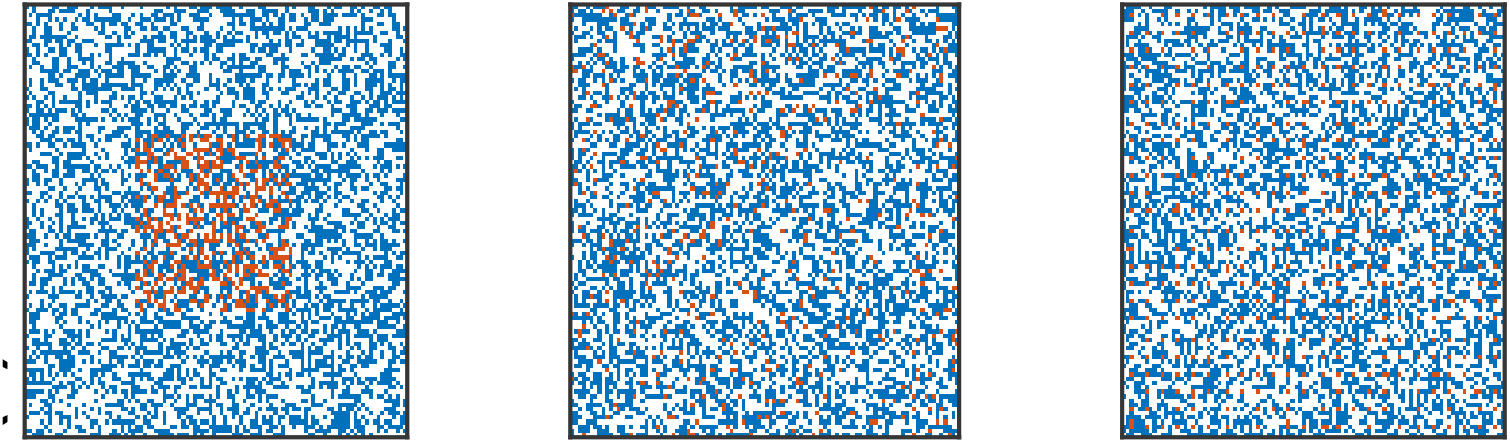
Initial cell configurations (clumped, random, and uniform, respectively). The red, blue, and white dots are the R-cells, S-cells, and empty sites, respectively.

### 2.2. Cell-Cycle Decision

Each cell occupies a lattice point in a square domain of size *l* × *l*. In every time step, each cell may stay stationary, or it can move, divide, or die. The S-cells and R-cells divide at rate *r*_*S*_ and *r*_*R*_, respectively. The S-cells and R-cells divide at a constant rate of *r*_*S*_ or *r*_*R*_, respectively. In this study, we considered the von Neumann neighborhood (VNHD), which comprised the sites on the east, west, south, and north of each cell. The death rate of both types of cells is *d*_*T*_. The drug concentration *D*(*t*) is homogeneous in the domain, and a drug-induced death rate (*δ*_*D*_) is applicable for S-cells only. A sensitive cell undergoing mitosis can be killed by a drug with a probability of *δ*_*D*_*D*(*t*). Both S-cells and R-cells follow the rules described in the flow chart in Figure 2. A brief explanation of the flow chart (Figure 2) is provided in the following.

**Figure 2.**
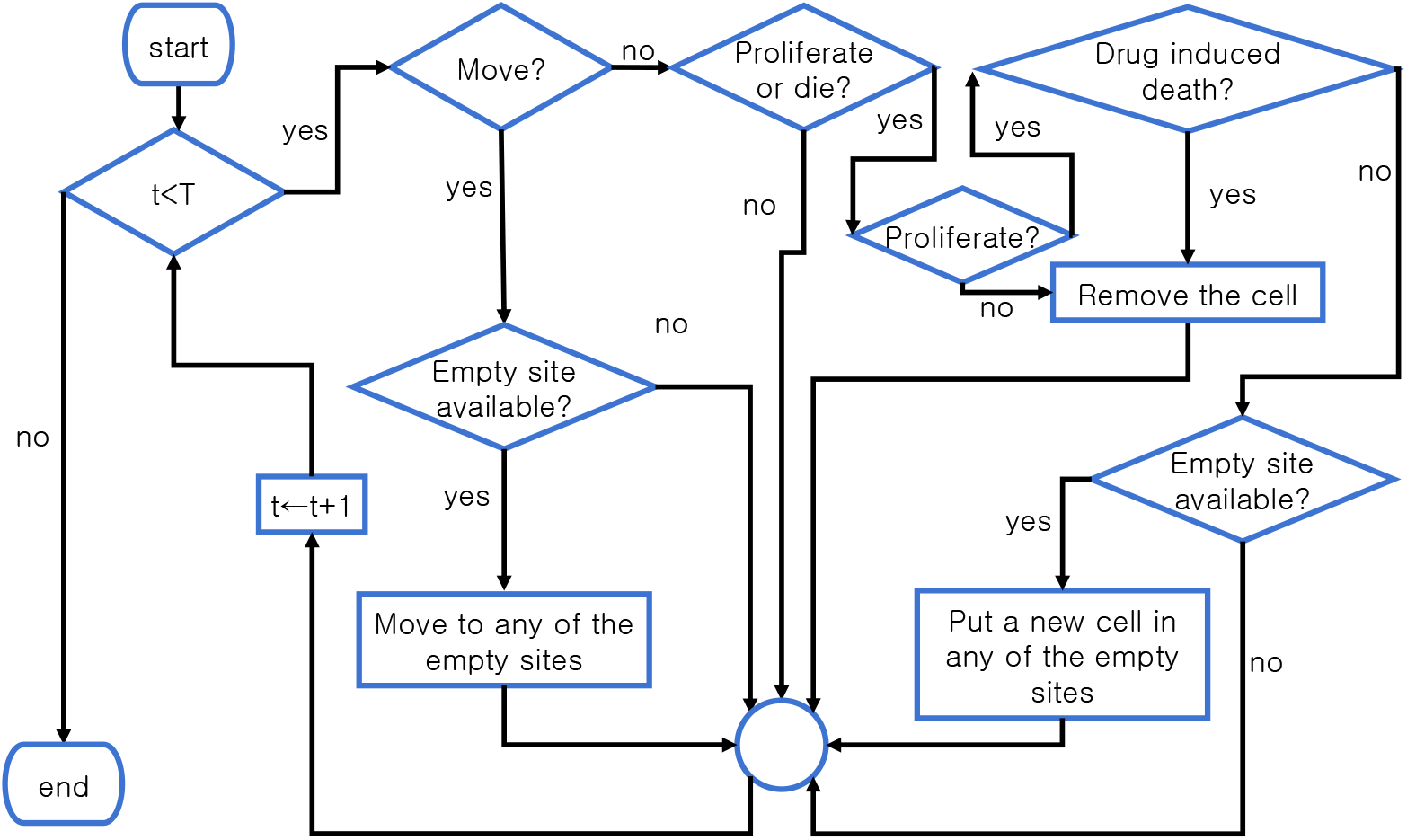
Flow chart of the cells’ life cycle. In each time step, all of the cells follow the steps in the flow chart.

- Step 1: If *t* < *T*, where *T* is the end time, go to Step 2; otherwise, go to Step 12.
- Step 2: Decide whether the cell will move. Pick a random number from a uniform distribution (*x*_*m*_ ∼ U[0, 1]). If *x*_*m*_ < *m*, where *m* is the probability of cell migration, then go to Step 3. If not, go to Step 5.
- Step 3: Is one of its VNHDs empty? If yes, go to Step 4. If not, go to Step 11.
- Step 4: Randomy move the cell to one of the empty sites in the VNHD. Go to Step 11.
- Step 5: Decide whether the cell will divide or die. Pick a random number from a uniform distribution (*x*_*pd*_ ∼ U[0, 1]). If *x*_*pd*_ < *r*_*j*_ + *d*_*T*_ with *j* ∈ {*S, R*}, where *r*_*j*_ is the j-cell proliferation rate and *d*_*T*_ is the normal cell death rate, then go to Step 6. If not, go to Step 11.
- Step 6: Decide whether the cell will divide. Pick a random number from a uniform distribution (*x*_*p*_ *∼* U[0, 1]). If 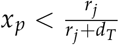, then go to Step 7. If not, go to Step 8.
- Step 7: Decide whether the cell will die due to the drug. If (*x*_*d*_ ∼ U[0, 1]) and if *x* < *δ*_*D*_*D*(*t*), where *δ*_*D*_ is the probability of cell death (for R-cells, *δ*_*D*_ = 0), go to Step 8. If not, go to Step 9.
- Step 8: Remove the cell, make the site empty, and go to Step 11.
- Step 9: Is one of its VNHDs empty? If yes, go to Step 10. If not, go to Step 11.
- Step 10: Randomly put a new cell of the same type in VNHD. Go to Step 11.
- Step11: *t* ← *t* + 1. Go to Step 1.
- Step12: The simulation ends.

### 2.3. Number of cells in the neighborhood

To quantify local cell–cell competition, we introduce the following notation.

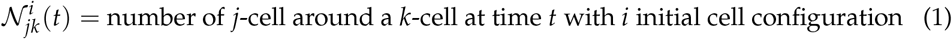

where *j* ∈ {*R*(R-cells), *S*(S-cells), *E*(Empty site)}, *k* ∈ {*R, S*} and *i* ∈ {*c*(Clumped), *r*(Random), *u*(Uniform)}. To denote the mean over all of the *k*-cells in the domain at time *t*, we write 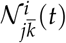.

We denote the number of empty sites by 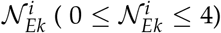. A cell can move to any unoccupied sites in its VNHD provided that 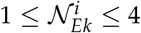. During cell proliferation, one parent cell divides into two daughter cells of the same type. To accommodate the daughter cell, at least one empty site is required 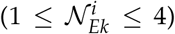 in the parent cell’s von Neumann neighborhood. If 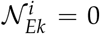, the proliferation was not executed. Upon cell division, one daughter cell is placed in the parent cell’s location, and the other is randomly placed in one of the empty sites in the VNHD. Upon the availability of an empty site in the VNHD (i.e., 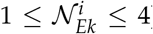), while attempting to divide, the mother S-cell may die with a probability of *d*_*D*_ due to the drug, but the R-cells do not experience drug-induced death. Dead cells are immediately removed from the respective sites.

### 2.4. Model Parameters

As a representative structure, we assume a square domain of 100 × 100 lattice points. We start our simulation with a tumor of *N*(0) = 5, 000 cells. The S-cells are assumed to be randomly dispersed over the domain. We assume that *f*_*R*_ = 10% of the cells are resistant. In the clumped case, all of the R-cells are randomly dispersed in a 40 × 40 clump. The parameters are summarized in Table 1.

**Table 1.** The parameter values are listed in the following table.

| Parameter | Description | Value | Reference |
| --- | --- | --- | --- |
| $t$ | Time in days | | |
| $T$ | Simulation end time | | |
| $S(t)$ | Number of sensitive cells | | |
| $R(t)$ | Number of resistant cells | | |
| $N(t)$ | Total number of cells | | |
| $K$ | Carrying capacity of each lattice point | 1, 2 | Assumed |
| $f_R$ | Initial percentage of the resistant cell population ($R(0)/N(0)$) | 10% | [36] |
| $r_S$ | Sensitive cell proliferation rate | 0.027 per day | [16] |
| $r_R$ | Resistant cell proliferation rate | $(1-0.3)r_S$ | [19] |
| $d_T$ | Cell death rate | $0.3r_S$ | [22] |
| $m$ | Migration rate | 0 to 100% of $r_S$ | Assumed |
| $D(t)$ | Drug concentration at time $t$ | 0, 1 | |
| $\delta_D$ | Drug-induced death rate of S-cells | 0.75 | [23] |
| $\rho$ | AT threshold for treatment break | 0.5 | [16] |
| $r_{SF}$ | Fibroblast-mediated sensitive cell proliferation rate | 200% of $r_S$ | Assumed |
| $r_{RF}$ | Fibroblast-mediated resistant cell proliferation rate | 200% of $r_R$ | Assumed |

### 2.5. Treatment Schedules

We consider two treatment strategies: continuous therapy (CT) and adaptive therapy (AT). In CT, the maximum tolerated dose (MTD) is applied to the domain over the entire simulation time. On the other hand, in AT, the treatment is provided from the beginning of the simulation until the cell population is reduced to *ρN*(0). The treatment is stopped until the total population, *N*(*t*), reaches *N*(0) again. Then, the treatment is re-applied. In this study, we assume that the MTD is applied during the treatment cycle and that *ρ* = 0.5 [16]. In mathematical notation, drug concentration can be written as follows. We consider the time when the total population reaches 120% of *N*(0) as the time to tumor progression (TTP).

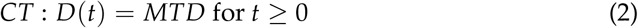

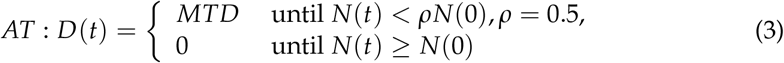

### 2.6. Simulation

The model was implemented on the JAVA platform using the Hybrid Automata Library (HAL) [37]. The generation of the initial cell configuration, the data analysis, and the visualization were performed by using MATLAB. To keep the results unbiased, the sequence of cells in the simulation was shuffled at the beginning of every time step. For each simulation scenario, we simulated 30 virtual tumors (i.e., 30 realizations of the model simulation), unless otherwise noted. To denote the average over the 30 simulations, we used over-bars, such as 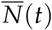 and 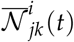.

### 2.7. Statistical Analysis and Progression Probability

To investigate the consequences of a parameter change in the results of the 30 realizations, we used a two-sample t-test. Significant differences with *p* − *value* < 0.001, 0.01, and 0.05 are represented by * * *, * * and *, respectively. For non-significant differences, we use “n.s.”. We also used a Kaplan–Meier survival curve to illustrate the approximate progression probability (PPr), which is defined as the relative frequency of tumor relapses among the 30 virtual tumors. Our results show an approximation of the PPr with respect to TTP. The Kaplan–Meier survival analysis was performed using MATLAB.

## 3. Results

### 3.1. Impact of the initial R-cell configuration on the time to progression under continuous therapy

First, we simulated CT on the three types of initial cell configurations for a time span of 2000 days. In this simulation, we assumed the carrying capacity of each lattice point to be *K* = 1 and the cell migration rate to be *m* = 0. Under CT, the S-cells died out quickly, and the remaining R-cells started to grow and fill the model domain. The representative spatial distributions of the tumor cells are shown in Figure 3(a) under CT on the 1^*st*^, 120^*th*^, and 2000^*th*^ day. The cell configuration in the clumped case was significantly different from those in the random and uniform cases, between which the difference seemed to be negligible. On the 120^*th*^ day, slightly larger patches of resistant cells are observed in the random case than in the uniform case. By the end of the simulation, the whole domain was captured by R-cells in both cases. On the other hand, in the clumped case, the R-cells grew in a patch in the center. By the end of the 2000 days, a huge clump of R-cells captured almost the entire domain.

**Figure 3.**
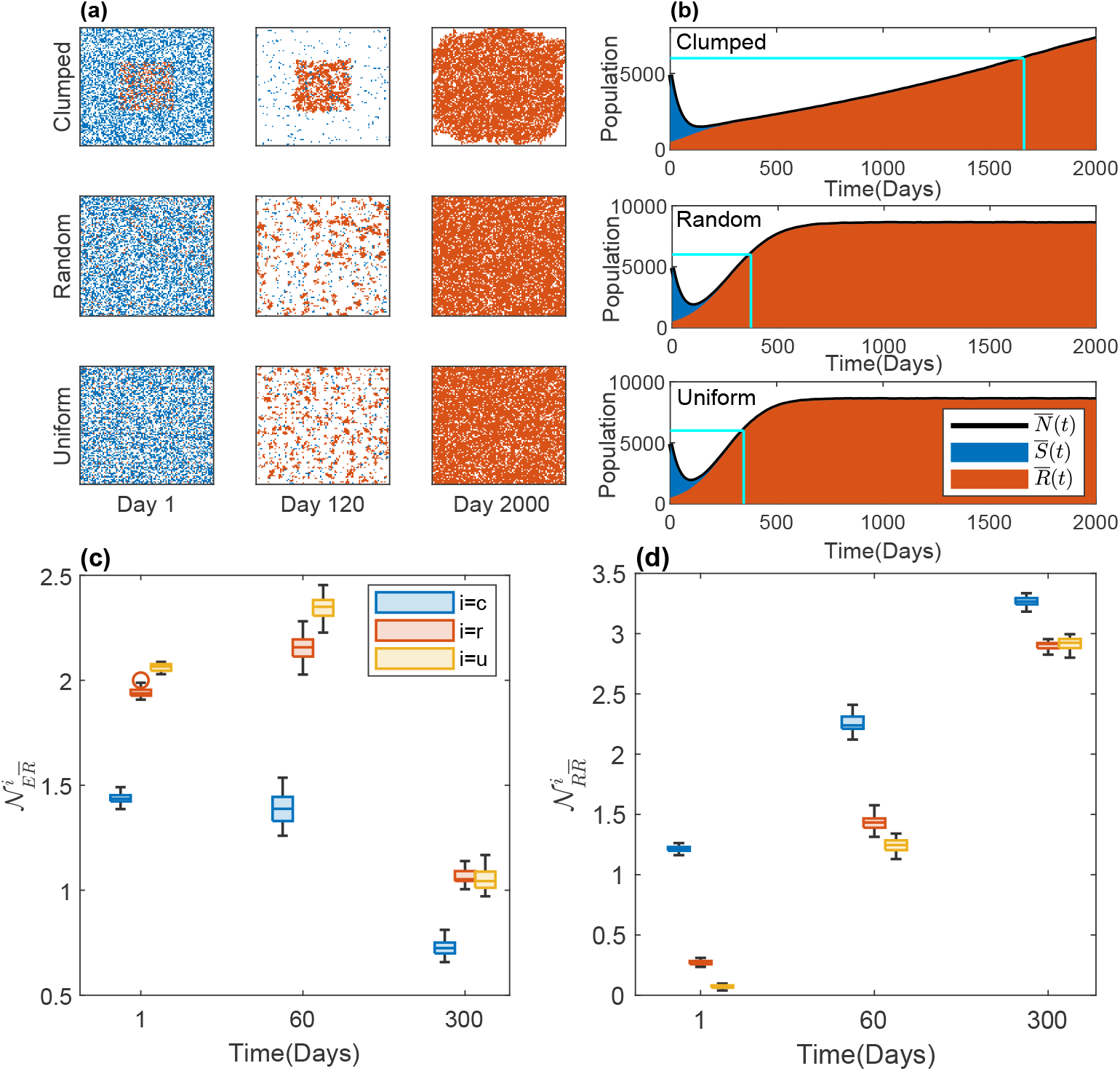
Effects of initial R-cell distribution on the TTP under CT. (**a**) The cell configurations on days 1, 120, and 2000 are shown. The blue, red, and white dots are S-cells, R-cells, and empty sites, respectively. (**b**) The average temporal evolution of the average number of S-cell and R-cell populations over 30 realizations with clumped (upper panel), random (middle panel), and uniform (bottom panel) initial cell configurations (blue: S-cell, red: R-cell). Black solid line: average total cell population 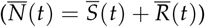. Vertical cyan line: TTP in each case; horizontal cyan line: the 120% level of the initial tumor volume (tumor progression threshold). The average numbers of empty sites 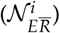 and R-cells 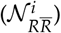 in the VNHD of an R-cell in the 30 realizations are shown as boxplots in (**c**) and (**d**), respectively, for *i* = *c, r, u*. The blue, red, and yellow boxes are for the clumped (*c*), random (*r*), and uniform (*u*) cases, respectively.

The temporal dynamics of different types of cells are presented in Figure 3(b). The TTP values in the three cases were 1662, 372, and 345, respectively. In Figure 3(b), the cyan horizontal line shows the 120% level of the initial tumor volume. The dynamics of the S-cells were almost same for all three types of initial configurations (Figure A1(a)), as they were initially similarly sparse and had the same growth parameters in all cases. Although the S-cell population dynamics were similar in all three cases, they affected the total cell growth by modulating the R-cells’ dynamics differently. To examine the reason for why the TTP was significantly different in the clumped case compared to the random and uniform cases, we investigated the local R–S and R–R spatial competition. Specifically, we calculated the numbers of S cells, R cells, and empty sites of each R-cell VNHD in the three spatial patterns.

To compare the local growth potential of R-cells in the three spatial patterns, we calculated the number of empty sites in the VNHD of each R-cell. The average number of empty sites in the neighborhood of an R-cell was lower in the clumped case than in both the random and uniform cases 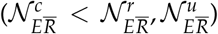 (Figure 3(c)). Figure 3(c) shows that the number of empty sites around each R-cell increased from day 1 to day 60 in the random and uniform cases (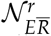 and 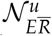 increased from day 1 to 60) because the treatment-induced deaths of S-cells freed up space in the neighborhood of each R-cell, leading to a reduction in R–S spatial competition. In the clumped case, however, the neighborhoods of the R-cells were mostly occupied with R-cells (Figure 3(a)), and thus, the number of empty sites in the VNHD of each R-cell 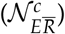 did not significantly increase (Figure 3(c)), blue boxes) after drug administration.

We next compared the R–R spatial competition by quantifying the average number of R-cells in the VNHD of each R-cell. In the clumped case, the number of R-cells in the neighborhood of each R-cell was significantly higher than in the random or uniform case 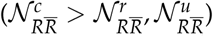 (Figure 3(d); blue boxes vs. yellow and orange boxes). The R–R competition increased over time in all of the cases; by the 300th day, about three R-cells were located in the neighborhood of each R cell, competing for a space to divide. Thus, each R-cell in the clumped case experienced, on average, greater spatial competition with R cells, leading to slow tumor progression. Taken together, our simulations demonstrate that the initial distribution of R-cells can modulate the time to progression under continuous therapy.

### 3.2. Impact of the initial R-cell configuration on the time to progression under adaptive therapy

Next, we investigated the effect of the initial R-cell distribution on the AT responses. Figure 4(a) shows a representative cell configuration at different times for AT with three different initial R-cell distributions. The cell population growth presented in Figure 4(b) shows that the TTP values were 1776, 392, and 362 days in the clumped, random, and uniform cases, respectively (vertical cyan lines). The total cell population went through four on–off treatment cycles until the TTP in the clumped case. In the other two cases, only one on–off treatment cycle was allowed until the TTP. To understand the mechanism by which AT caused a more delayed TTP in the clumped case compared to the random or uniform case, we first investigated the local growth potential on days 1, 35, and 95. We chose the 35th day (when the first cycle had yet to finish) and the 95th day (after which the second cycle started) for all the cases. The average numbers of empty sites (4(c)) and S-cells (4(d)) in the neighborhood were lower in the clumped case than in the other two cases 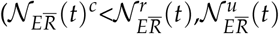 (Figure 4(c)). The number of empty sites in the neighborhood of a each R-cell 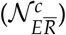 did not significantly change from day 1 to day 35, though the numbers in both the random and uniform cases (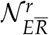 and 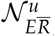) increased remarkably. During the first treatment break (from day 35 to day 95), the S-cells divided, filling up empty sites in the neighborhoods. This resulted in a reduction of 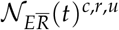 (Figure 4(c): boxplots on day 95 vs. boxplots on day 35).

**Figure 4.**
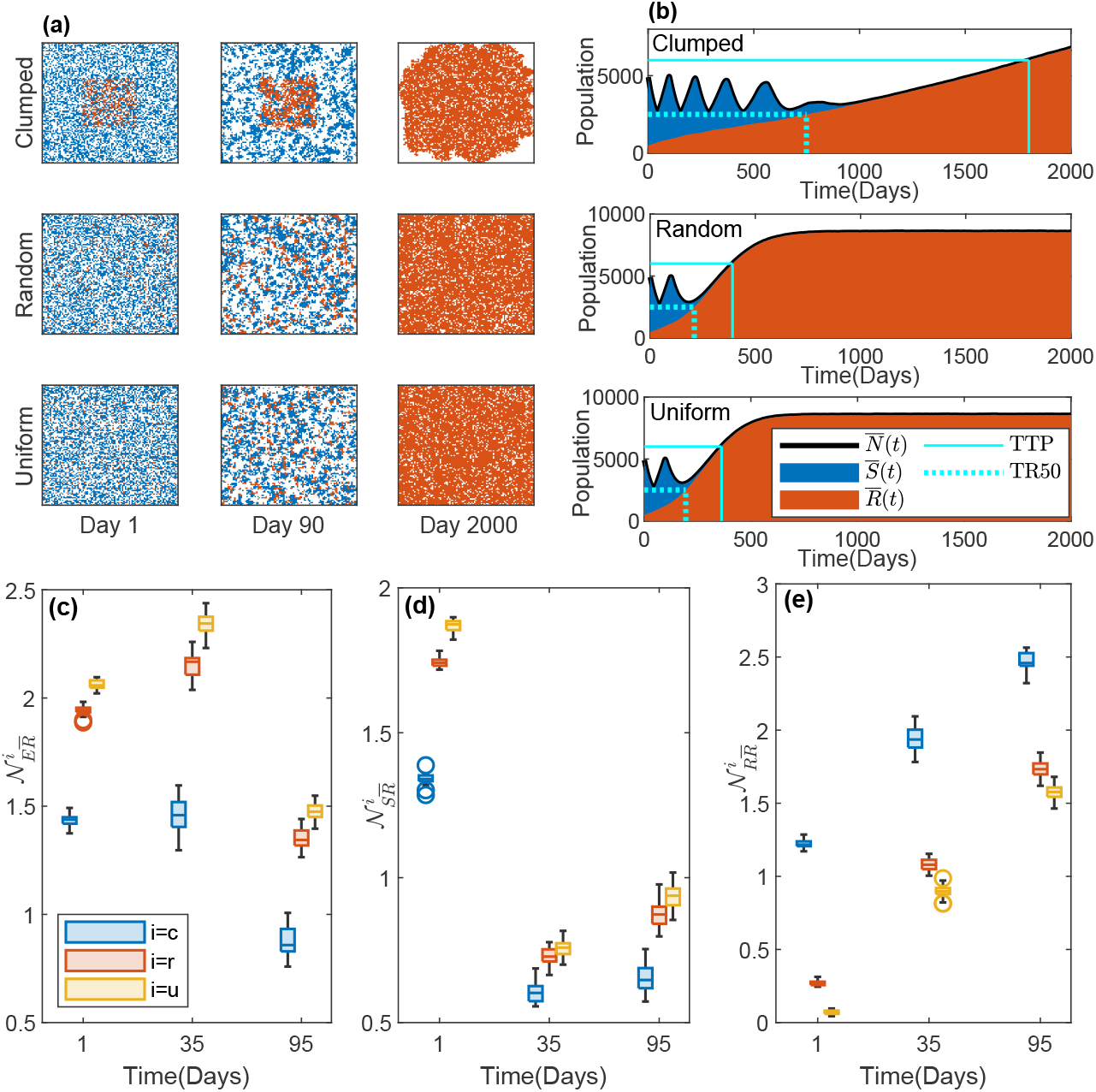
Effect of the initial R-cell distribution on the time to progression under AT. (**a**) Cell configurations on days 1, 120, and 2000. The square signifies the domain representative of the tumor. The blue, red, and white dots are S-cells, R-cells, and empty sites, respectively. (**b**) Average temporal evolution of the S-cell and R-cell populations for the clumped (upper panel), random (middle panel), and uniform (bottom panel) cases of the initial cell configurations (blue dots: S-cells, red dots: R-cells). Black solid line: the total population (*N*(*t*) = *S*(*t*) + *R*(*t*)). Vertical solid cyan line: TTP; horizontal solid cyan line: 120% level of the initial tumor volume. Vertical dotted cyan line: time for the R-cells to reach 50% of the initial tumor volume (TR50); horizontal dotted cyan line: 50% level of the initial tumor volume. The average numbers of empty 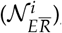, S-cells 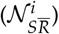, and R-cells 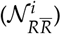 in the VNHD of an R-cell in the 30 realizations are shown as boxplots in (**c**), (**d**), and (**e**), respectively, for *i* = *c, r, u*. The blue, red, and yellow boxes are for the clumped (*c*), random (*r*), and uniform (*u*) cases, respectively.

Next, we compared the intensity of the spatial competition between the S-cells and R-cells. The average number of S-cells in each R-cell neighborhood was higher in the random and uniform cases than in the clumped case 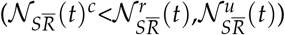 (Figure 4(d): yellow/orange boxplots vs. blue boxplots). The difference between the average number of S-cells in a neighborhood in the clumped case and those in the other two cases decreased from the 1st day to the 35th day (i,e., 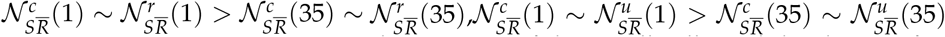). The higher number of S-cells in the VNHDs of the R-cells allowed the drug to free up sites more in the random and uniform cases than in the clumped case. In the first treatment break (from day 35 to day 95), the number of S-cells in the neighborhoods increased in all three cases due to the proliferation of S-cells during the “off” part of the treatment cycle (Figure 4(d)). Thus, the inhibition of growth of R-cells by S-cells was higher in the random and uniform cases than in the clumped case.

Finally, we quantified the strength of inter-species spatial competition (i.e., competition between R-cells). The average number of R-cells in a neighborhood in the clumped case was always higher than the numbers in the random and uniform cases 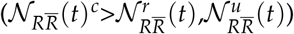 (Figure 4(e)). Interestingly, the number of R-cells in each R-cell neighborhood in all three cases increased irrespective of drug administration because R-cells can proliferate regardless of drug administration. A greater number of R-cells in the neighborhood implies a stronger inhibition of R-cell growth by R-cells, leading to a slower rate of cell population growth (Figure 4b: slope of the total population growth in the clumped case < slope of the total population growth in the random and uniform cases) .

In summary, in the random and uniform cases, the the number of R-cells increased more quickly due to the space available in the neighborhoods, and it reached the level of 50% of the initial total cell population during the end of the second drug administration after the first “off” part of the treatment cycle (Figure 4b: dotted cyan line). Once the number of R-cells reached the level of 50% of the initial cell population, the ongoing (additional) cycle of treatment could not reduce the total population below the 50% level, leading to continuous treatment and a quicker progression (Figure 4b). Under CT, inter-species competition (R–R competition) was solely responsible for determining the TTP (higher competition leading to delayed TTP). Under AT, however, a combination of R–R competition and R–S competition seemed to determine the TTP. In other words, a more significant reduction in the growth inhibition of S-cells by R-cells combined with the increase in R–R competition drove a faster TTP in the random and uniform cases. In the clumped case, the R–R competition is the main determining factor of TTP.

### 3.3. Clumped initial distribution results in higher clinical time gain (TG)

So far, we explored the impact of the initial R cell distribution on the AT and CT outcomes and investigated how the treatments modulate the inter- and intra-species competition (R–S and R–R, respectively), resulting in different treatment outcomes. During CT, the drug is supplied consistently without considering the response, which causes a prompt decline in the S-cell population and facilitates R-cell growth by lowering the local R–S spatial competition. On the other hand, during AT, the drug is supplied in short cycles to keep a tolerable number of S-cells, which are required in order to limit the R-cells’ growth by maintaining spatial competition. Therefore, AT is expected to maintain total cell growth for longer than CT. We quantified the benefits of AT over CT in terms of the TG (= TTP in AT - TTP in CT). The TG is a consequence of spatial competition, which depends on the distribution of R-cells and their neighborhoods’ occupancies. Our results show that TG was significantly higher in the clumped case than in the other two cases (Figure 5(a), p-value < 0.001). Furthermore, an even higher TG could be observed if the initial fraction of R-cells was smaller (Figure 5(b) *f*_0_ = 1%). In Figure 5 (a) and (b), the random and uniform cases do not show significant differences.

**Figure 5.**
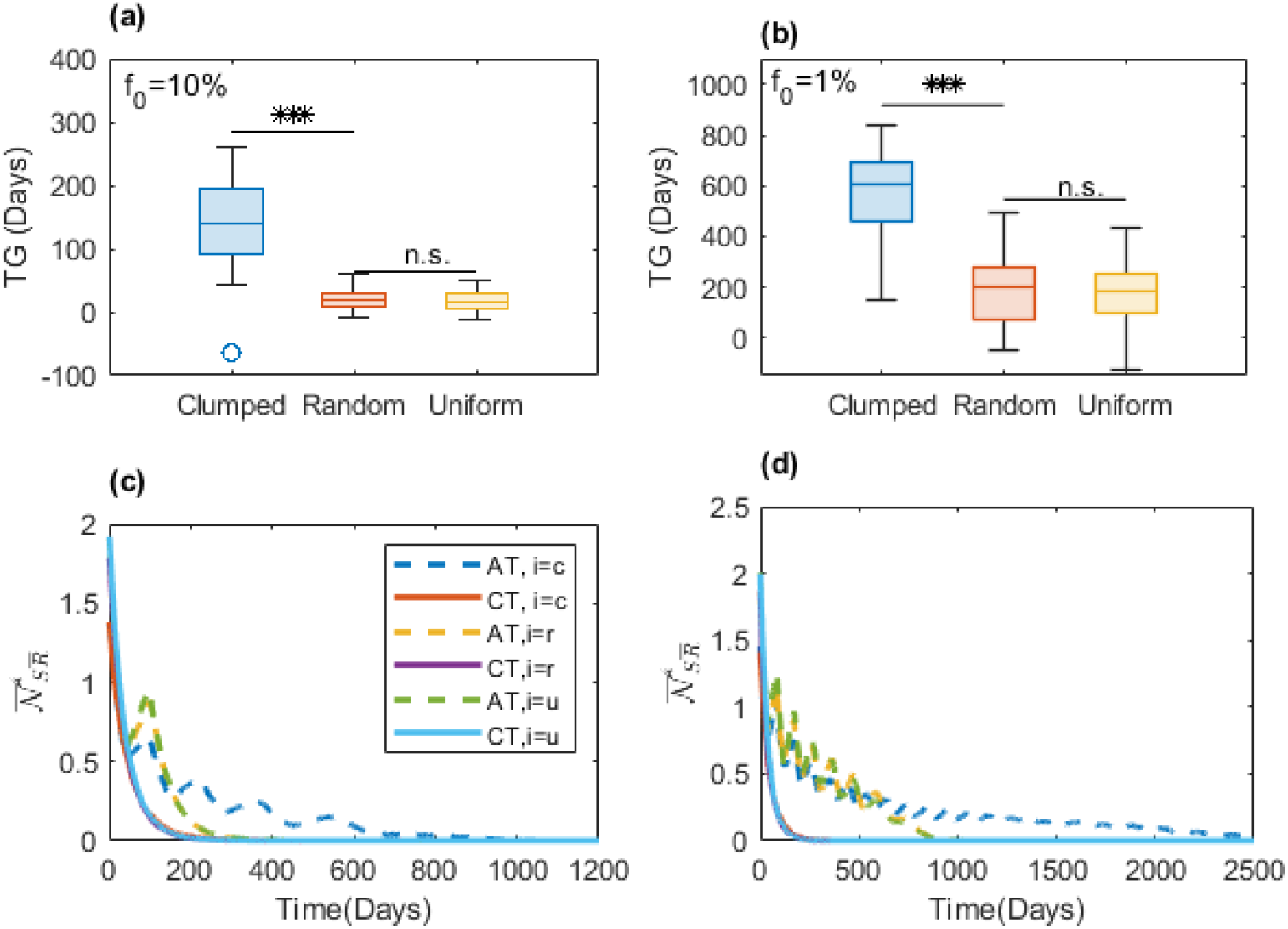
Role of the initial R-cell distribution on the benefit of AT over CT. (**a**) The boxplot shows the TG in the 30 realizations for the clumped, random, and uniform initial cell configurations for *f*_0_ = 10% (* * *: *p* − *value* < 0.001; n.s.: not significant). (**b**) Similar results to those in (a) are shown for *f*_0_ = 1%. (**c**) The time evolution of the mean of the average number of S-cells in the VNHD of an R-cell in the 30 realizations 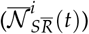 is shown for both CT (solid lines) and AT (dashed lines); c: clumped, r: random, u: uniform. The solid lines overlap, but the blue dashed line (AT, i = c) shows the longer existence of S-cells in the VNHD of an R-cell in the case of the clumped initial cell configuration under AT. (**d**) Similar results to those in (c) are shown for *f*_0_ = 1%.

To understand the role of sustained S-cells, we plotted the mean of 30 realizations of the average number of S-cells in each R-cell’s neighborhood 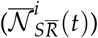 against time. Figure 5 (c–d) shows that in the clumped case, S-cells are maintained for longer in the neighborhood than in the random and uniform cases. This longer existence of the S-cells in the clumped case allowed the AT therapy to significantly increase the TG compared to the TG achieved with CT. Furthermore, we observed that there was a slight chance of attaining a negative TG. For the clumped, random, and uniform cases, this was 0.03, 0.13, and 0.17, respectively, for *f*_0_ = 10%, or 0, 0.07, and 0.1 for *f*_0_ = 1%. This shows that a clumped distribution of resistant cells increases the preference for AT over CT.

The uniform and random cases did not show significant differences in terms of TG. Furthermore, the clumped case and random case were two extreme versions of similar types of distributions. Thus, we focused on the case of clumped R-cell distribution in our further simulations.

### 3.4. Increased carrying capacity reduces the benefit of adaptive therapy by reducing spatial competition

For the clumped initial cell distribution, we investigated the effect of the spatial carrying capacity on the TG. The spatial carrying capacity was characterized as *K* = 1 (each lattice point could hold one cell) or *K* = 2 (each lattice point could hold, at most, two cells, regardless of their sensitivity or resistance). When *K* = 1 was used, a total of four cells could occupy the VNHD of each cell (i.e., 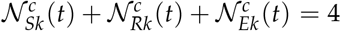). For each cell in *K* = 2, a total of eight cells could occupy a VNHD, and one additional cell could be located in the respective cell’s site (i.e., 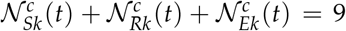). Figure 6(a) shows that increasing the carrying capacity significantly decreased the TG (*p* − *value* < 0.001) from a median of 139 days to a median of 7 days. Increasing the carrying capacity provided additional room for accommodation of the daughter cells, which is observed in Figure 6(b). Initially, the number of empty sites in each R-cell 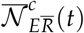 was above 5 for *K* = 2, whereas it was below 2 for *K* = 1. Due to this ample space in their neighborhoods, R-cells hardly experienced any spatial competition and grew at a higher pace when *K* = 2 under both AT and CT. As the total cell population grew, 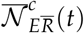 decreased abruptly and tended to settle below 1. For *K* = 1, a similar trend was observed; however, the number of empty sites was lower than that for *K* = 2 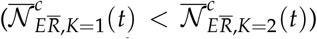 .Comparing the number of empty sites in each R-cell’s VNHD 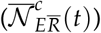 for AT in the case of *K* = 1 with that in the case of *K* = 2 (Figure 6(b), dotted lines)), we observed that, for 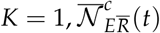 went through ups and downs several times, which suggested spatial competition with neighboring cells. On the other hand, for *K* = 2, this value monotonically decreased, and there was a very slight difference due to AT and CT. Therefore, we concluded that the short TG with *K* = 2 was due to the lack of spatial competition. We observed that the probabilities of having a negative TG were 0.03 and 0.4 for *K* = 1 and 2, respectively, i.e., an increase in carrying capacity reduces the benefit of AT over CT.

**Figure 6.**
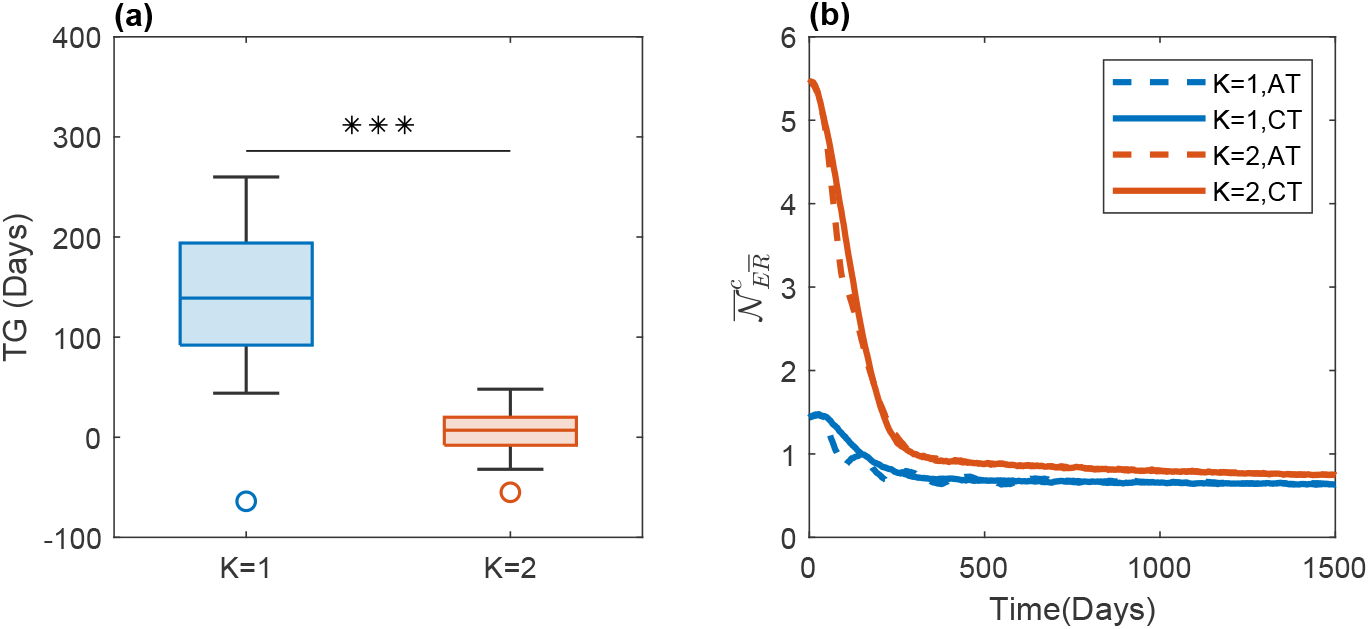
Effect of carrying capacity on the time gain (TG). (**a**) The blue and red boxplots show the TG from the 30 couple realizations (for both AT and CT) with respect to carrying capacities of *K* = 1 and 2, respectively. The triple asterisk (* * *) signifies that increasing the carrying capacity significantly reduced the TG (*p − value* < 0.001). (**b**) The time evolution of the mean of the average number of empty sites in the VNHD of each R-cell in the 30 realizations 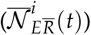 is shown for both CT (solid lines) and AT (dashed lines); *K* = 1 (blue) and 2 (red). *K* = 2 offers a greater number of empty sites in the VNHDs of R-cells than *K* = 1.

### 3.5. Increased cell migration rate reduces the time gain of AT compared to CT

To investigate the impact of cell migration on therapeutic responses, we simulated a model of the migration rate *m* = 0%, 50%, and 100% of the birth rate (of the respective types of cells (Table 1)). Under both AT and CT, an increased cell migration rate promoted faster cell population growth, leading to a shorter TTP (Figure 7(a)). The time to progression without cell migration was 1667 under CT. The TTP decreased to 1282 when the cell migration rate increased to 50%, and further increased to 1075 when the rate was 100%. On the other hand, the time to progression without cell migration was 1803 under AT. The TTP decreased to 1376 when the cell migration rate increased to 50%, and further increased to 1152 when the rate was 100%.

**Figure 7.**
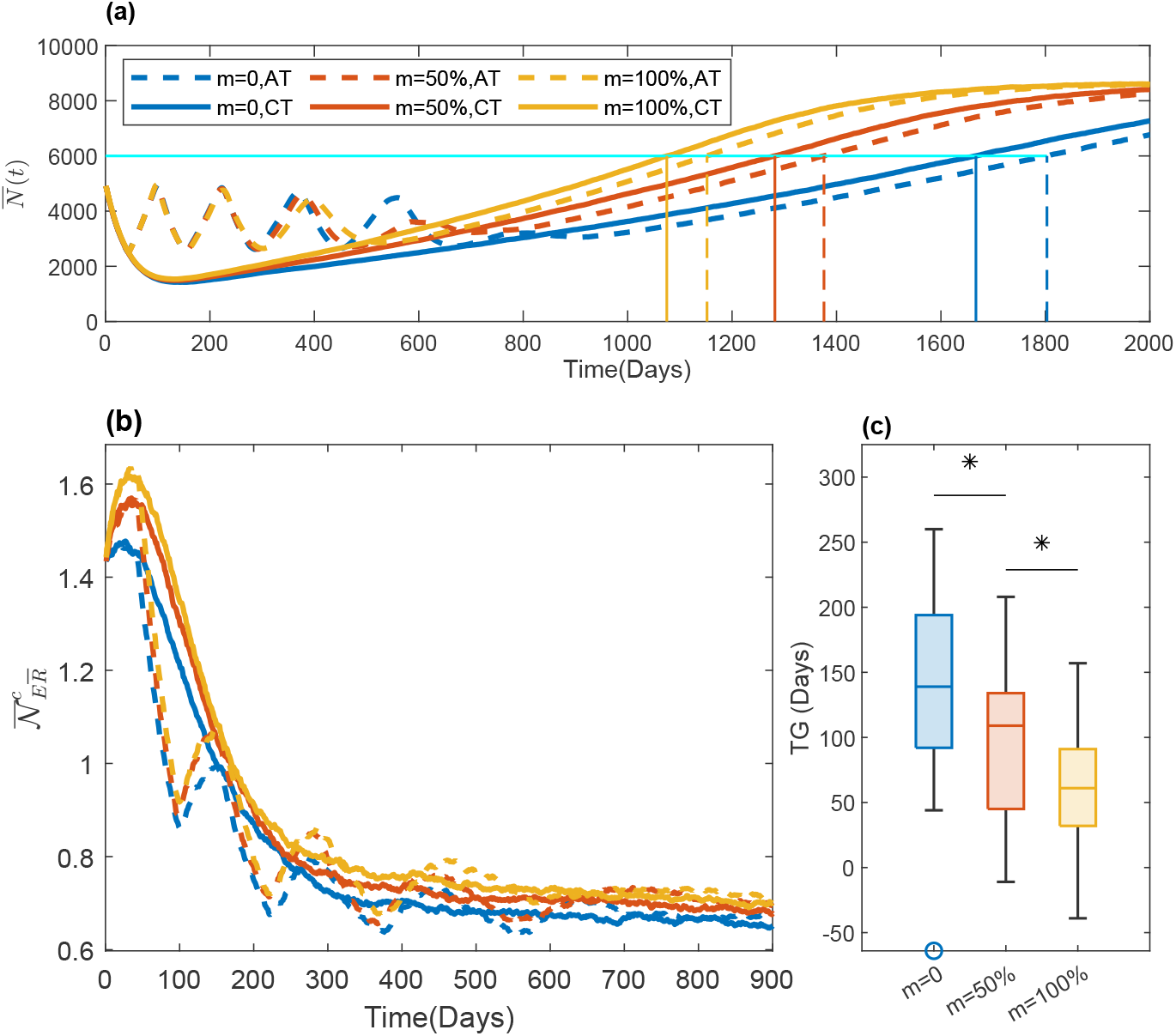
Effects of cell migration on TG. (**a**) The time evolution of the mean of the total cell population 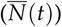 in the 30 simulations is shown for *m* = 0 (blue), 50% (red), and 100% (yellow) under AT (dashed line) and CT (solid line). The vertical solid lines show the time to tumor progression (TTP) under CT. The vertical dashed line shows the TTP under AT. (**b**) The time evolution of the mean of the average number of empty sites in the VNHD of each R-cell in the 30 realizations 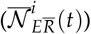 is shown with the same line styles as in (a). (**c**) The blue, red, and yellow boxes show the boxplots of the TG for *m* = 0%, 50%, and 100%, respectively. * : p-value < 0.05.

To understand the mechanism by which migration causes a faster relapse, we investigated the temporal evolution of the local growth capacity (number of empty sites in the VNHD of each R-cell; 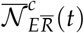) (Figure 7(b)). The figure shows that 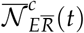 was smaller for *m* = 0 than for *m* = 50% and 100% under CT. We observed a similar impact of cell migration on the AT response. During the “on” period of the treatment, the S-cells died, resulting in an increase in the number of empty sites in the VNHD of each R-cell in all of the migration rate cases. The lowest increase, however, was observed in the absence of migration. This increase in empty sites in each R-cell’s neighborhood as a result of cell migration implies a higher growth potential for each R-cell, leading to a faster treatment failure. Therefore, cell migration reduces the local spatial R–R competition, leading to a rapid increase in the total cell population. Furthermore, the average number of empty sites is lower under AT than under CT; i.e., competition is higher under AT. Therefore, migration relaxes competition and increases the growth rate under AT more than under CT, which results in a significant reduction in the time gain due to migration (*p − value* < 0.05).

### 3.6. Fibroblast-mediated drug resistance

So far, we investigated the role of initial R-cell distribution, carrying capacity, and cell migration on the therapeutic response. Fibroblasts are known to promote cancer cell growth and drug resistance [30–34]. In particular, a recent paper by Marusky et al. revealed the impact of fibroblast location on the outcomes of continuous therapy [6]. To understand the impact of fibroblast distribution on the outcomes of adaptive therapy, we considered three types of fibroblast configurations: (i) in the absence of fibroblasts (NoF), (ii) in the presence of fibroblasts that overlap with clumped R-cells at the center (FC), and (iii) in the presence of fibroblasts around the initial clump of R-cells (FSq). In the case of FC, we assumed the existence of a fibroblast as a 32 × 32 square at the center, and in the case of FSq, we considered the fibroblast region as a hollow square with an outer dimension of 55 and inner dimension of 45, diagonally situated between the sites (23, 23) and (77, 77), with a wall thickness of five lattices. Figure A2 shows a graphical illustration of the three fibroblast configurations. We assumed that fibroblasts comprised about 10% of the domain. Although fibroblasts are known to promote cancer cell growth at varying rates [33,34], for simplicity, we assumed that fibroblasts could increase the growth of both R- and S-cells by 200% (doubling their respective cell growth rates: *r*_*SF*_ = 200% of *r*_*S*_ and *r*_*RF*_ = 200% of *r*_*R*_) [33]. Under CT, the total cell population decrease until about day 125, 144, or 176 in the cases of FSq, NoF, and FC, respectively (Figure 8). Though it is decreasing, the less stiff red line shows a smaller reduction rate in the case of FC than for the other two configurations (blue (NoF) and yellow (FSq)). For FC, the locations of all of the R-cells initially belong to the locations of the fibroblast (Figure A2: the fourth row). Thus, the growth of all R-cells is promoted by the fibroblasts. In the other configurations, none of the R-cells initially belonged to the fibroblast region and did not gain fibroblast-mediated increases in their growth rates. The net growth rate of R-cells for FC was higher than that in the other two cases. As a result, at the minima, the total cell population in the case of FC remained the highest among the three types of configurations (1416, 1423, and 1681 cells for FSq, NoF, and FC respectively). Once all of the sensitive cells are eradicated by CT and the clump of the resistant cells was saturated with cells, the growth dynamics of the total cell population in NoF and FC became similar because the elevated growth rate due to the presence of the fibroblast at the center had no impact, spatial competition near the center was already higher (Figure A2: column 3). On the other hand, in the case of FSq, R-cells grew remarkably faster than in the other two cases when the R-cells reached the fibroblast region (Figure A2: columns 2 and 3, row 6). From about the 500th day onward, the cells on the circumference of the clump exceeded the fibroblast region, and the fibroblast region became saturated with R-cells (Figure A2: column 4, row 6). Thus, all of the configurations exhibited similar growth rates, which are represented by the parallel-looking growth curves (Figure 8(a); day 500 onward). Boxplots of the corresponding TTP values accompanied by the Kaplan–Meier plots of the PPr are shown in Figure 8(b) for the 30 realizations. The tumor progression in FSq was significantly faster than in both NoF and FC (*p* − *value* < 0.001).

**Figure 8.**
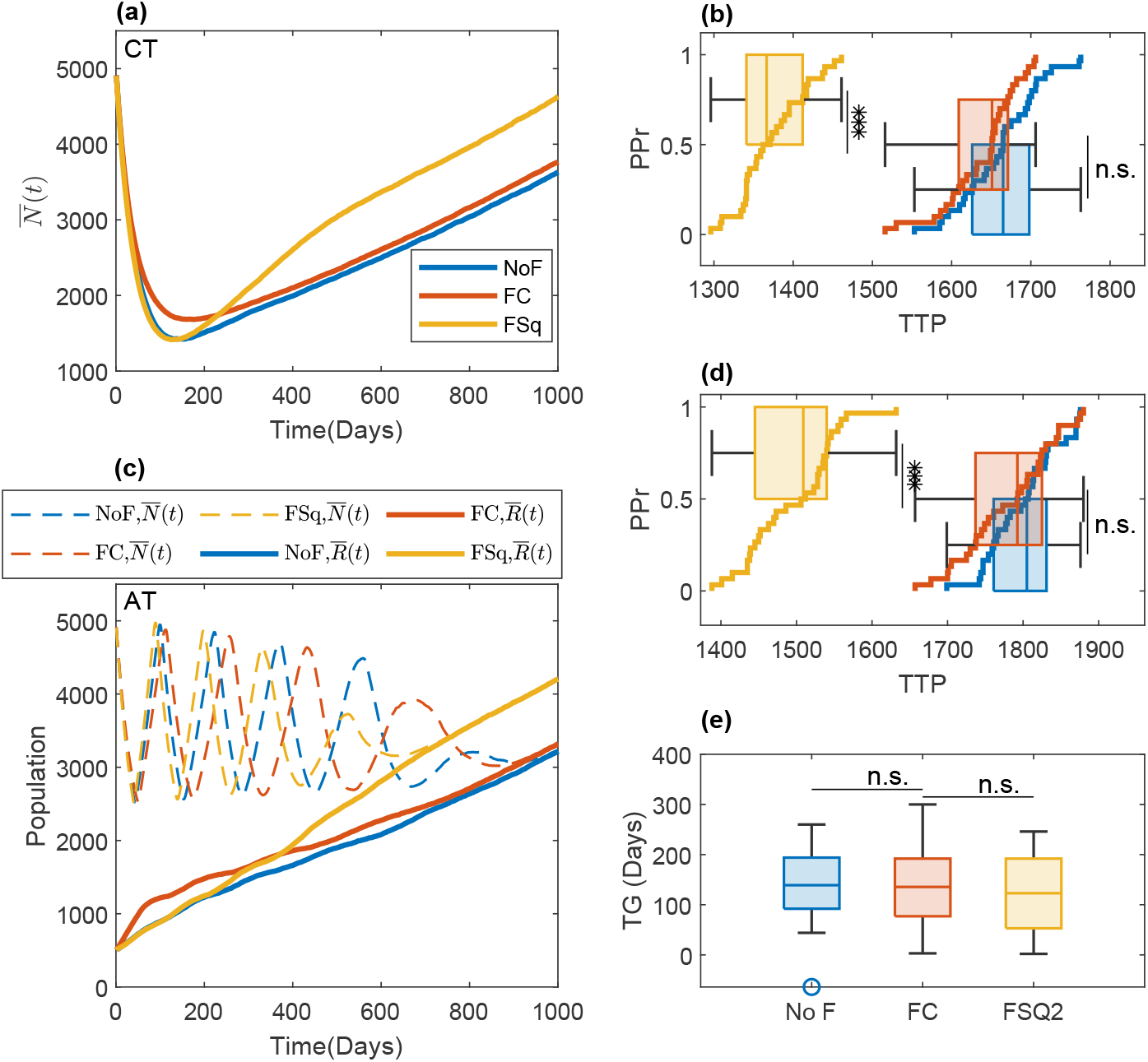
Consequences of fibroblast-mediated resistance in the clumped R-cell distribution. (**a**) The time evolution of the mean of the total cell population 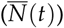 in the 30 simulations is shown for CT for three types of fibroblast configurations—no fibroblast (NoF), fibroblasts overlapping with clumped R-cells in the center (FC), and fibroblasts encapsulating the clumped R-cells in the center (FSq); these are shown with blue, red, and yellow lines, respectively. (**b**) Boxplot of the time to progression (TTP) in the 30 realizations under CT, along with the progression probability (PPr) for the three types of fibroblast configurations (NoF (blue), FC (red), and FSq (yellow)). (**c**) The time evolution of the mean of the total cell population 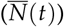, R-cells 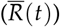, and S-cells 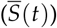 is shown with dashed, solid, and dotted lines for the 30 simulations under AT for the three types of fibroblast configurations (NoF, FC, and FSq—shown by blue, red, and yellow lines, respectively). (**d**) Boxplot of the time to progression (TTP) in the 30 realizations under AT, along with the progression probability (PPr) for the three types of fibroblast configurations (NoF (blue), FC (red), and FSq (yellow)). (**e**) The blue, red, and yellow boxplots show the time gain (TG) under AT compared to CT for the three types of fibroblast configurations (NoF, FC, and FSq, respectively). Though with the FSq configuration, a reduction in TG was observed under both CT (in (b)) and AT (in (d)), no significant differences were observed in the TG.

A similar type of scenario was observed for the growth of R-cells under AT. The growth of R-cells under AT with the clumped initial condition is shown in Figure 8(c) (solid lines) for the three different fibroblast configurations. During the first “on” treatment, R-cells initially grew the fastest with FC due to overlapping-fibroblast-mediated growth promotion (Figure A2). After about the 90th day, the growth rates for all of the fibroblast configurations were similar until about the 210th day. After the 210th day, the R-cells grew faster for FSq than for the other two configurations because, by this time, the R-cells reached the fibroblast region (Figure A2), and the R-cells on the circumferential area of the clump exploited the fibroblast-mediated growth, as well as the absence of spatial competition with other R-cells. The corresponding TTP values are shown in Figure 8(d) as boxplots, which are accompanied by Kaplan–Meier plots of the PPr. The disease progression was significantly faster in Fsq than in the other two cases. Interestingly, there were no significant differences in the benefits of AT (time gain; TG) because in our simulation, fibroblast-mediated protection of S-cells was negligible, while its promotion of R-cell growth was far more significant (Figure 8(e)).

We also simulated the effect of fibroblast location on the random initial R-cell distribution. We observed that both FC and FSq showed quicker progression than NoF under both CT and AT (Figure 9(a–d)). Although both the FC and FSq fibroblast configurations had almost the same surface area (i.e., the same number of sites with fibroblasts), FSq had a greater perimeter (both outer and inner) than FC. With FC, the R-cells with higher growth rates were closely located to the same kinds (i.e., fibroblasts promoted R-cells), leading to even a higher spatial competition between R-cells. In FSq, a good number of cells with higher growth rates competed with cells with normal growth rates. As a result, FSq elevated the net growth rate of more cells than FC although they had almost the same number of fibroblasts, leading to quicker progression than that in NoF and FC. The time to progression was significantly different in all cases (Figure 9(b,d), *p* − *value* < 0.001 in all cases). However, in terms of TG, we again observed no significant differences (Figure 9(e)).

**Figure 9.**
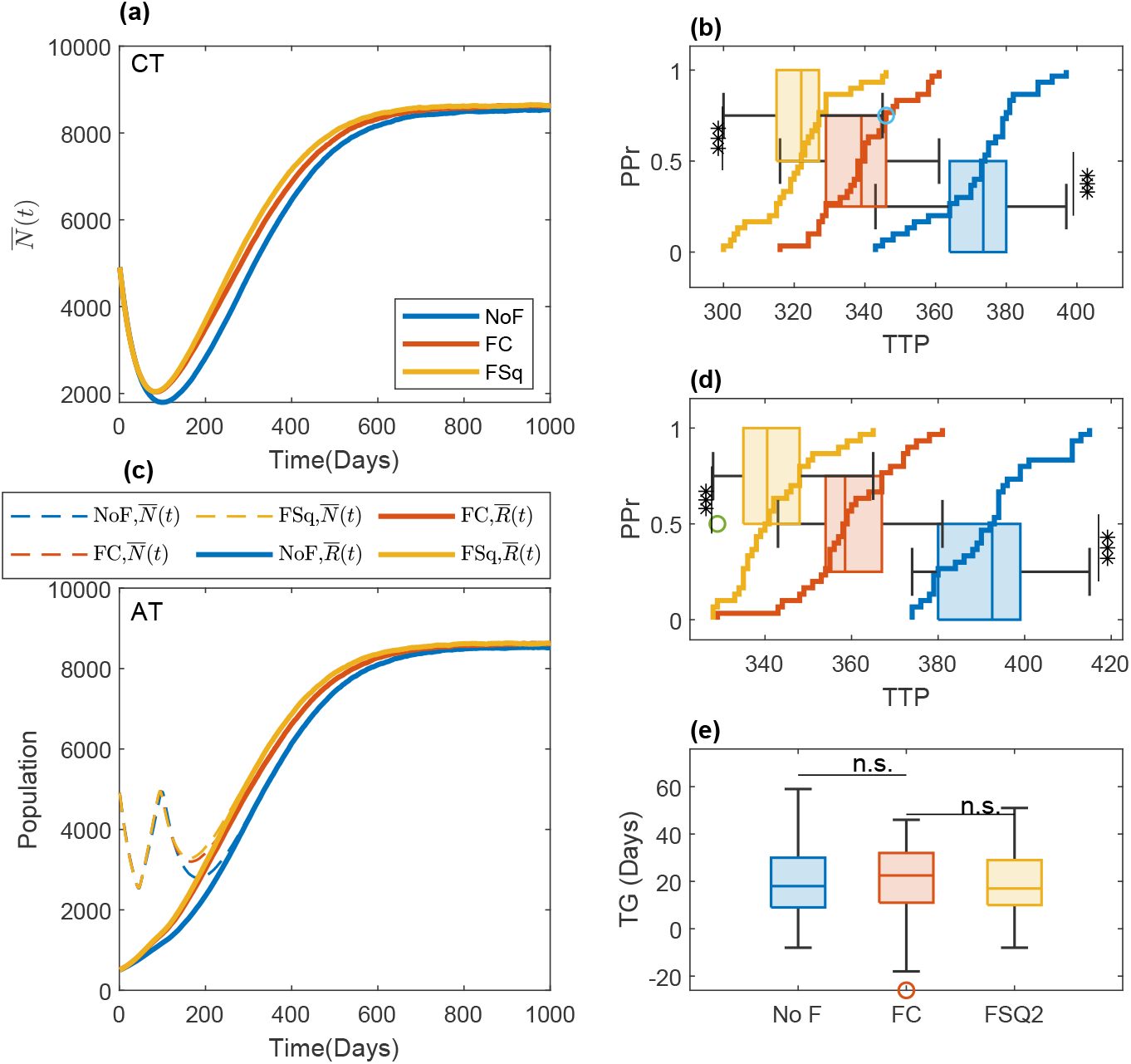
Effect of fibroblast location on tumor progression and the benefit of adaptive therapy compared to continuous therapy. (**a**) The time evolution of the average of the total cell population 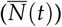 under CT in the 30 simulations is shown for the three types of fibroblast configurations—no-fibroblast (NoF), fibroblast center overlapping with clumped R-cells (FC), and fibroblast encapsulating clumped R-cells (FSq)—with blue, red, and yellow lines, respectively. (**b**) Boxplot of the time gain under CT in the 30 realizations, along with the progression probability (PPr) for the three types of fibroblast configurations: NoF (blue), FC (red), and FSq (yellow). (**c**) The temporal evolution of the averages of the total cell population 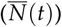, R-cells 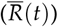, and S-cells 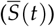under AT in the 30 simulations is shown with dashed, solid, and dotted lines for the three types of fibroblast configurations—NoF, FC, and FSq, which are represented by blue, red, and yellow lines, respectively. (**d**) Boxplot of the time gain under AT in the 30 realizations, along with the progression probability (PPr) for the three types of fibroblast configurations: NoF (blue), FC (red), and FSq (yellow). (**e**) The blue, red, and yellow box plots show the time gain under AT compared to CT for the three types of fibroblast configurations: NoF, FC, and FSq, respectively. No significant differences in the TG were observed.

To explore the impacts of both cell migration and fibroblast location on therapeutic outcomes, we additionally simulated the model with the clumped R-cell distribution and a cell migration rate of *m* = 50% (Figure A3). The result was similar to that of the case with the clumped initial distribution and *m* = 0% (Figure 8). However, as we observed before (Figure 7), due to the reduction in R–R competition as a result of cell migration, tumor progression occurred earlier for all types of fibroblast configurations and treatment strategies (AT and CT).

### 3.7. Adaptive therapy on a virtual patient with multiple metastatic lesions: three detected lesions and one undetected lesion at the beginning of therapy

In the sections above, we investigated the treatment response with a single tumor lesion (either a primary or metastatic site). Patients with advanced cancers who undergo the systematic therapy that we consider in this study typically present with multiple metastases. To understand the impact of the spatial heterogeneity of R-cells and fibroblasts on treatment outcomes, we simulated AT and CT in a virtual patient with four metastatic lesions, each of size 200 × 200 (we increased the metastasis size here to comply with the fibroblasts). Each metastatic lesion had its own independent domain, in which the cells were subject to space constraints. However, all metastatic lesions were subjected to the same systemic treatment, which was guided by a systematic biomarker that was represented by the total number of cells in all metastatic lesions. The characteristics of the local microenvironments were significantly different. For instance, the numbers of fibroblasts were different among the metastatic sites. Due to the different compositions and densities of extracellular matrixes, tumor cell migration can be different. We considered four combinations, which are shown in Figure 10, and considered a tumor consisting of four metastases that held the four different biological combinations. In addition, we assumed that one of the metastases was invisible (contained too few cells to be detected initially). We assumed that the number of tumor cells in the invisible metastatic site was 10% of number of cells in the other metastases. Therefore, we modeled four different cases of tumors with four metastases, of which one metastatic lesion was invisible (presented by the red color in Figure 10). For the visible metastases, we assumed a clumped initial R-cell distribution. We also assumed that the total number of cells was 10 times the number of R-cells (*N*(0) = 10*R*(0)) in each of the metastases. The S-cells were randomly distributed over each metastatic lesion’s domain. The locations of the fibroblasts were assumed to be scattered. We simulated these four metastatic tumor models for two types of invisible metastases: (i) All the tumor cells belonged to a 60 × 60 grid centered with the metastases (clumped), or (ii) all of the tumor cells were sparse over the all of the metastases (random). In both cases, the R-cells made up 10% of the total cell population and were dispersed randomly over the respective areas. Figures 11 and 12 show the initial cell configurations of the four metastatic lesions in the four cases mentioned above (in Figure 10) for the clumped and random invisible metastases.

**Figure 10.**
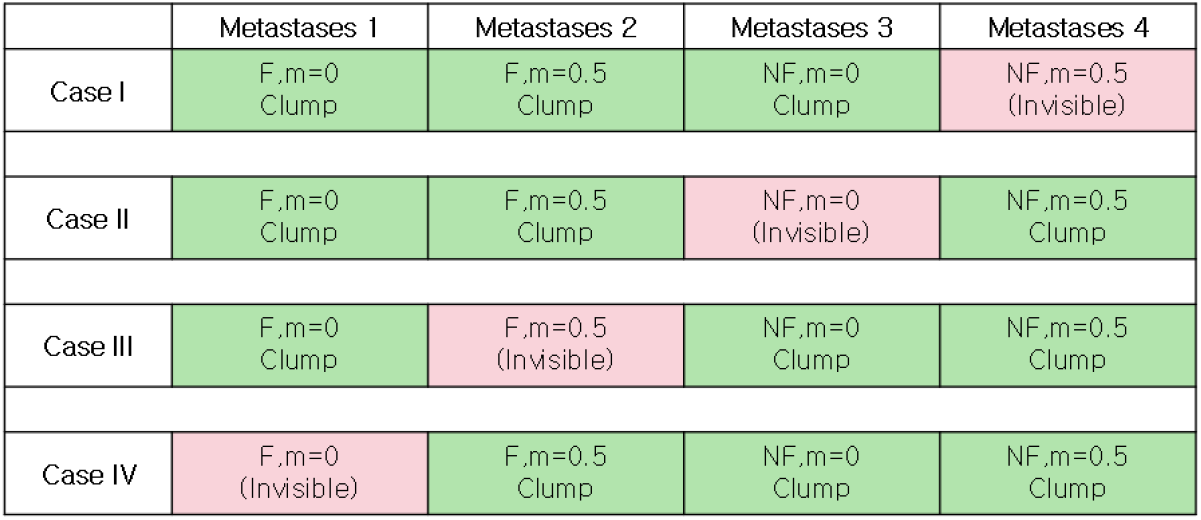
Combinations of the four metastasis scenarios. The green and red colors correspond to detected and undetected metastases, respectively. F and NF correspond to the existence and absence of fibroblasts, respectively.

**Figure 11.**
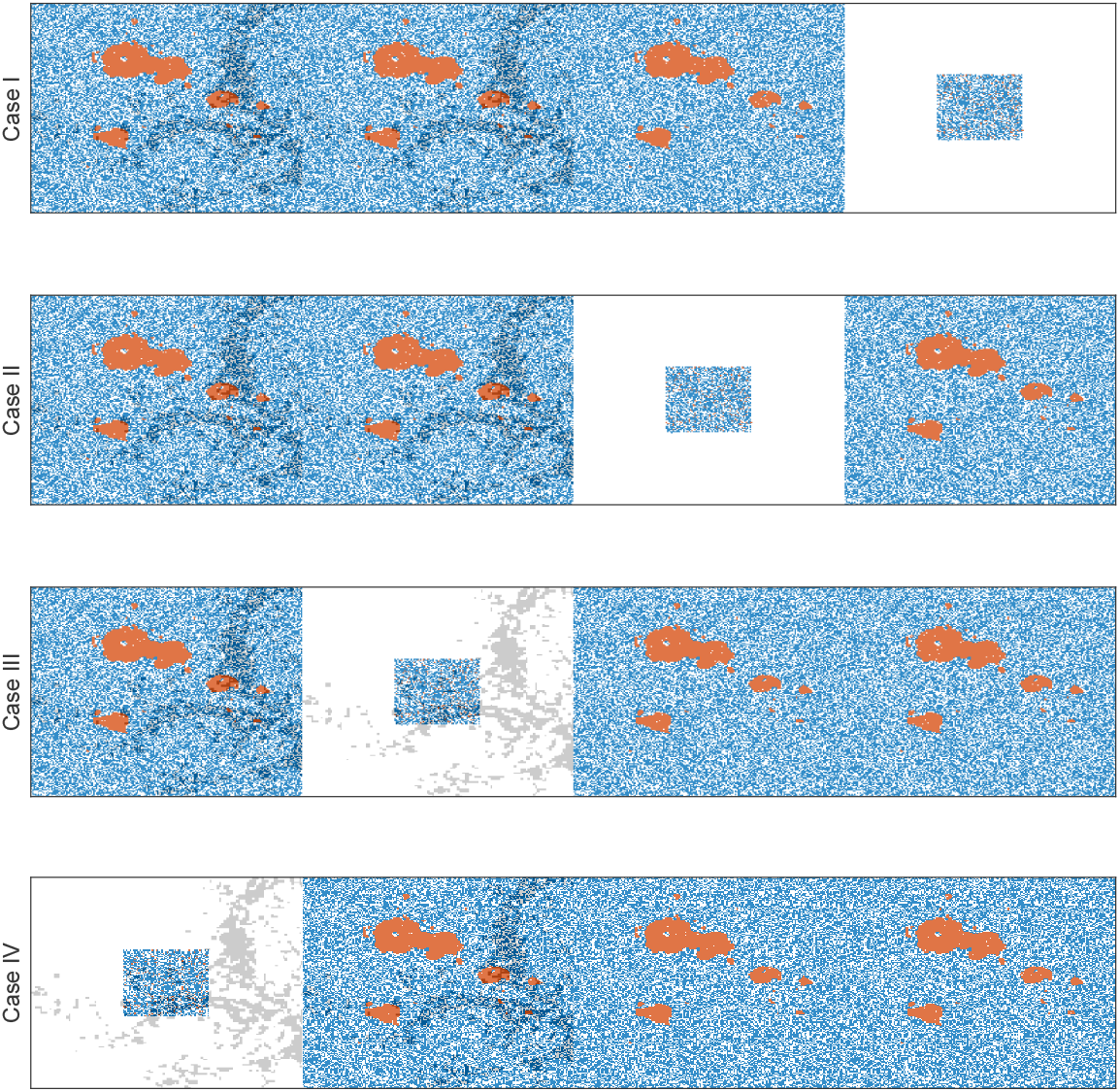
Initial cell configurations for the four cases with an invisible clumped metastasis. The red, blue, and white dots correspond to R-cells, S-cells, and empty sites. The gray dots shows the sites accompanied by fibroblasts.

**Figure 12.**
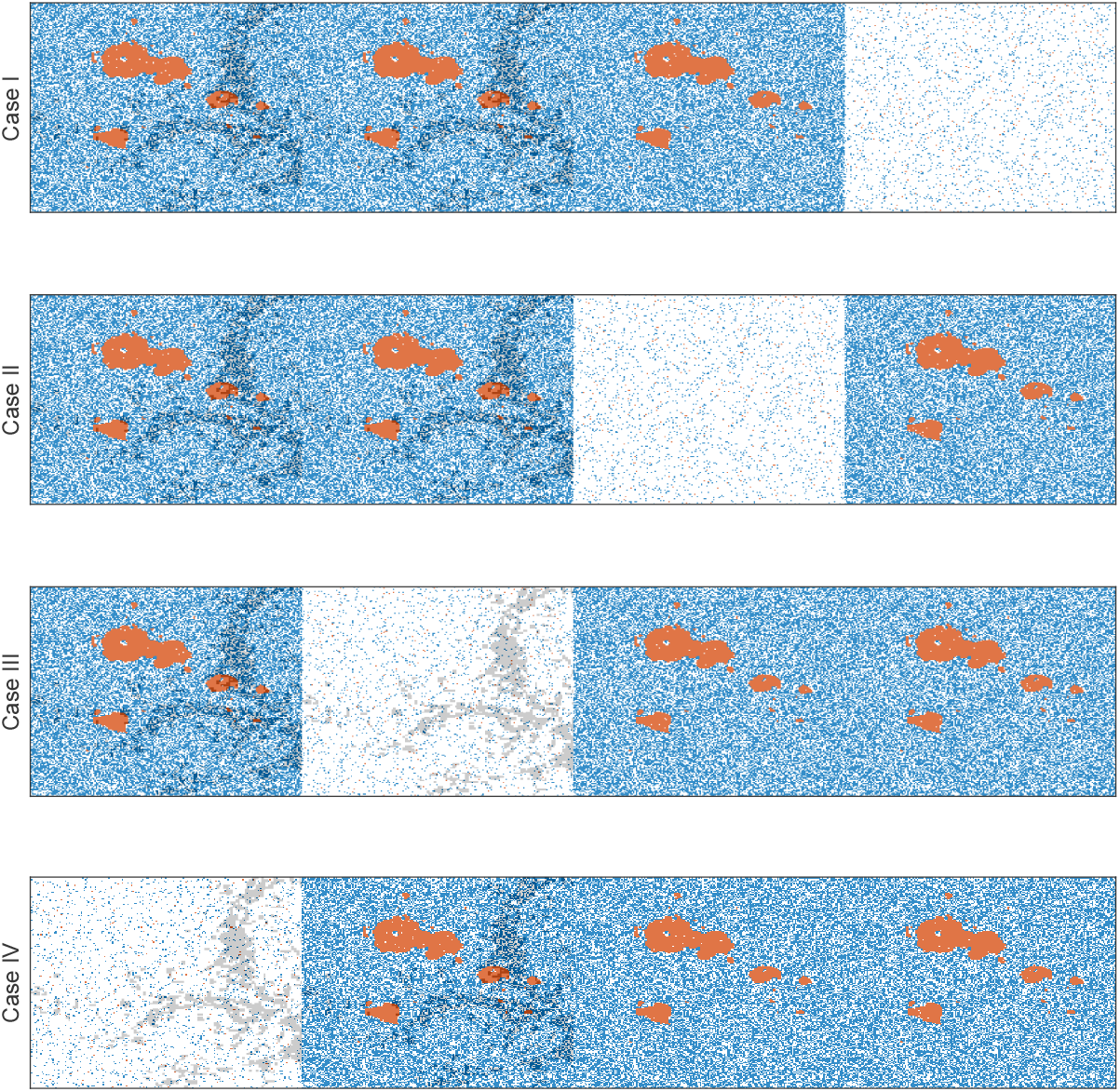
Initial cell configurations for the four cases with a random invisible metastasis. The red, blue, and white dots correspond to R-cells, S-cells, and empty sites. The gray dots show the sites that are accompanied by fibroblasts.

In these simulations, we use two different criteria for tumor progression: emergence time (ET) and TTP. ET was defined as the time for a new metastatic site to be detected, which was assumed to be the time when the total cell population was 50% of the overall domain’s carrying capacity in the respective metastasis. The TTP was defined as the time when the total cell population of the four metastases reached 120% of the total initial cell population.

We observed that both cell migration and fibroblasts can promote faster relapses (shorter TTP) in Sections 3.5 and 3.6. Similar consequences were observed here. For instance, in case I with the clumped invisible metastatic lesion (Figure 13, first graphs in the left column), the total cell population grew faster in metastatic lesion 2 than in metastatic lesion 1 (Figure 14) due to the higher cell migration probability in metastasis 2. The total cell population grew faster in metastasis 1 than in metastasis 3 (Figure 14) due to the fibroblasts in metastasis 1. The invisible metastasis (metastasis 4) become noticeable on day 2632 under CT and on day 2633 under AT (Figure 14, rows 2 and 3, respectively) when the total number of cells in the fourth lesion reached 50% of the domain carrying capacities of those specific metastases. The ET was almost the same for CT and AT (vertical solid cyan line vs. vertical dashed cyan line), but the TTP in CT was shorter than that in AT (solid (CT, 2076 days) and dashed (AT, 2302 days) red lines). The cell configurations are shown in the fourth and fifth rows of Figure 14. Most importantly, when tumor progression had already occurred, the invisible metastasis had not yet reached a detectable tumor size. We observed a similar order in the growth of the tumor cell population in metastases 1 to 3 in Case I, as well as with the random invisible metastasis (Figure 13, left vs. right figures). The cell configurations at crucial times are shown in Figure 15. The cell growth in the random invisible metastasis was much faster than in all other metastases, in agreement with the results in Sections 3.1 and 3.2. Importantly, the resistant cell populations in this metastatic lesion experienced less competition with the sensitive cell population because the duration of the systematic therapy determined by the sum of all metastatic lesions was so long that most sensitive cells in the lesion were killed off by the first cycles. The random distribution imposed less competition between the resistant cell populations, resulting in the rapid growth of resistant cells. The fourth invisible metastasis became the largest on day 399 under CT and on day 574 under AT.

**Figure 13.**
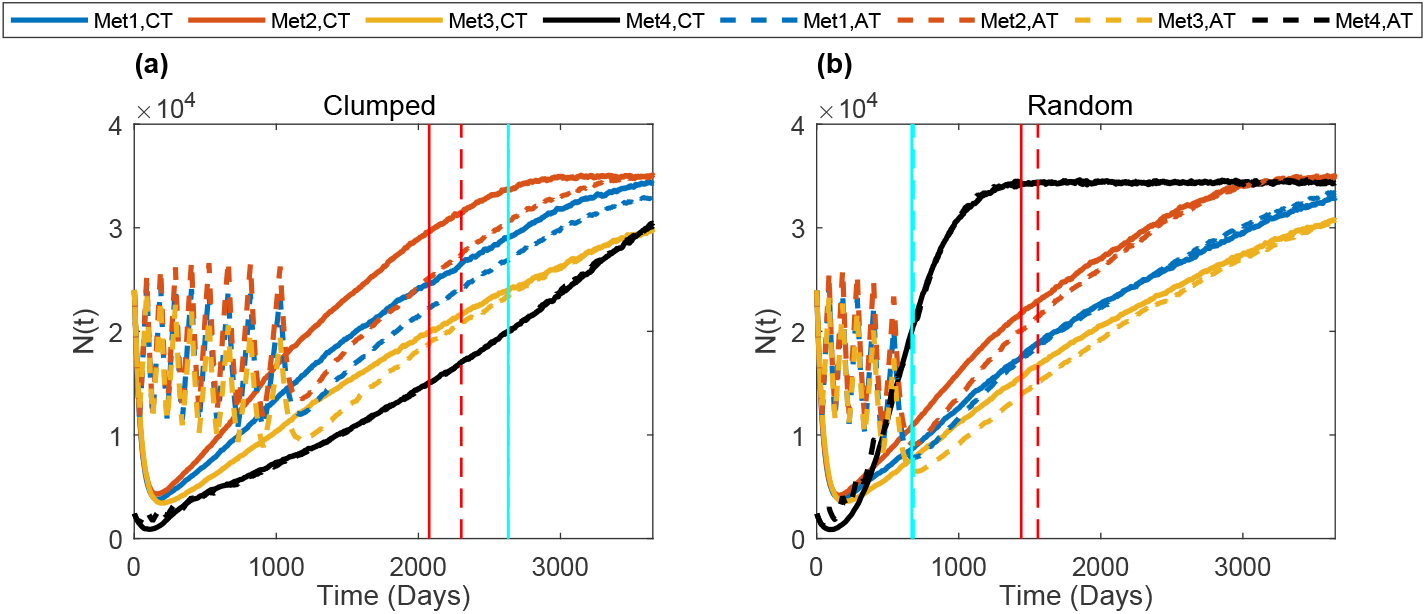
Complex dynamics of multiple metastases under AT and CT. The time evolution of the total cell population in the four metastases is shown in the sub-figures. The first, second, third, and fourth rows show the results for Case I. The first and second columns show the results for the initial clumped and random cell configurations in the invisible metastasis, respectively. In each sub-figure, the blue, red, yellow, and black colors show the total cell populations in metastasis 1, metastasis 2, metastasis 3, and metastasis 4, respectively; the vertical cyan lines show the emergence time (ET) of the invisible metastasis, and the red line shows the TTP. The solid and dashed lines show results under CT and AT, respectively.

**Figure 14.**
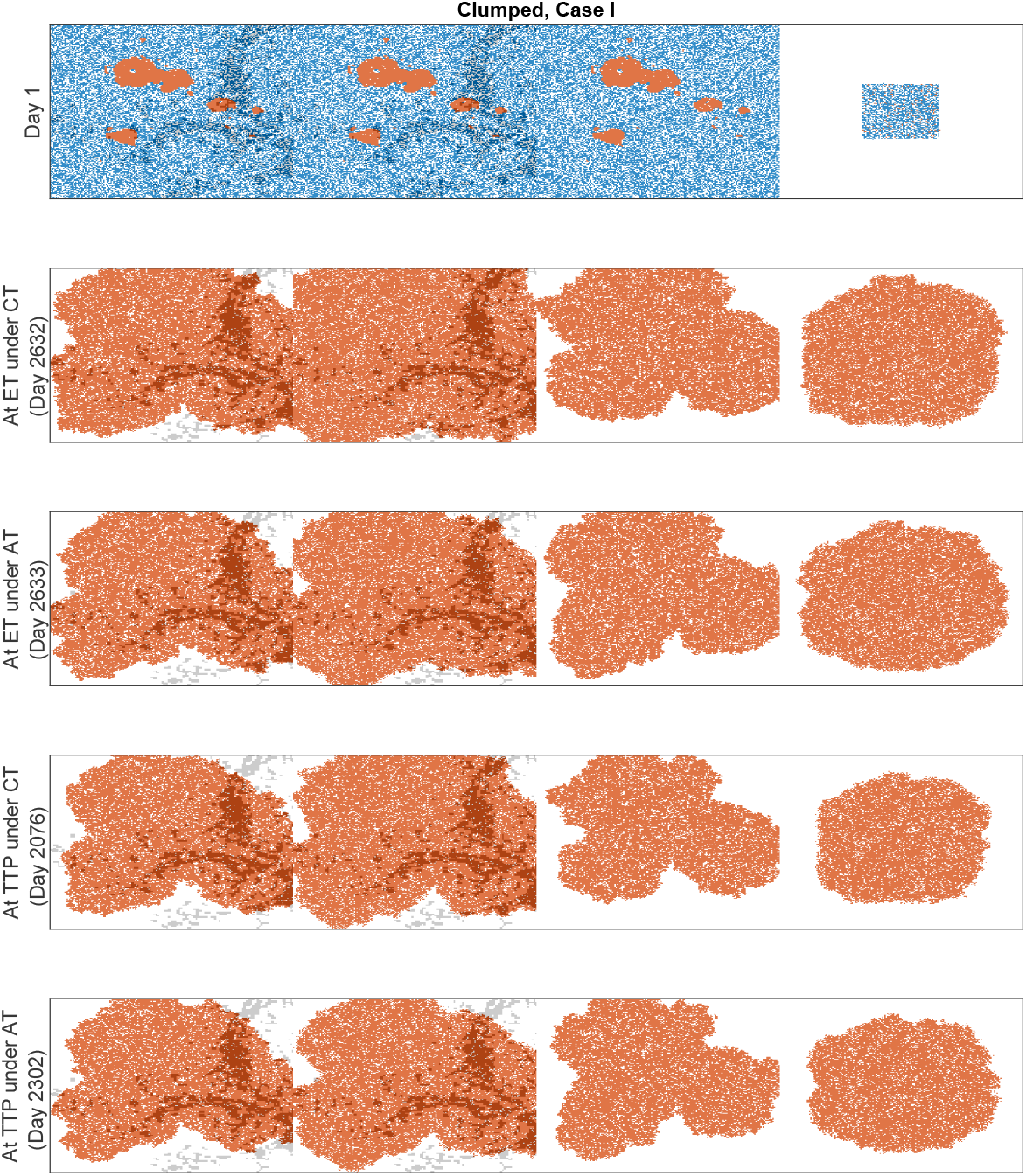
Cell configurations at the ET and TTP under CT and AT for Case I with the clumped invisible metastasis. The red, blue, and white dots correspond to R-cells, S-cells, and empty sites. The gray dots show the sites that are accompanied by fibroblasts.

**Figure 15.**
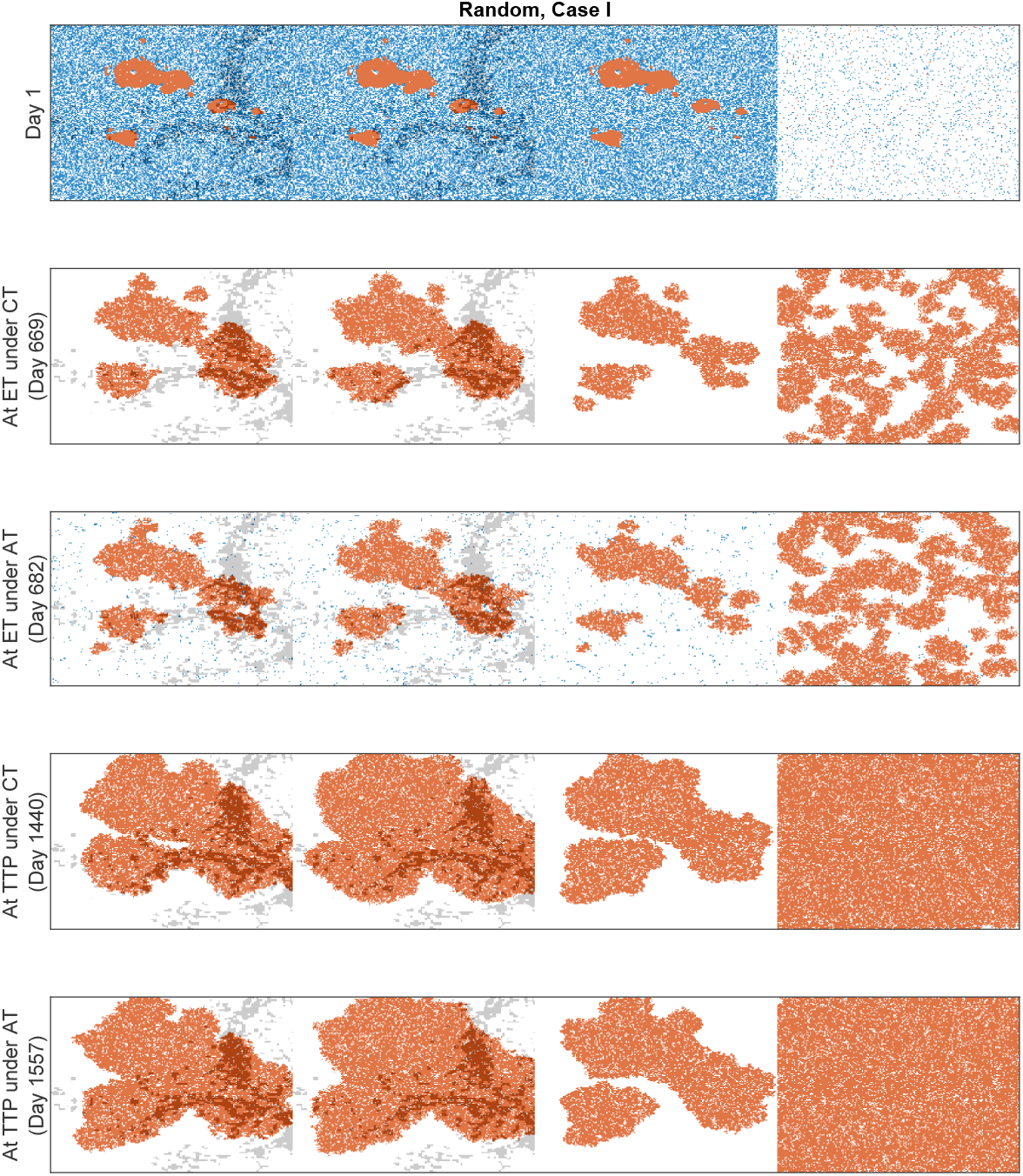
Cell configurations at the ET and TTP under CT and AT for Case I with the random invisible metastasis. The red, blue, and white dots correspond to R-cells, S-cells, and empty sites. The gray dots show the sites that are accompanied by fibroblasts.

For Cases II, III, and IV, similar results were obtained (Figure A4). A comparison of the ET and TTP in the four cases is shown in Figure 16 The ET was more delayed in Case II than in Case I, as the higher cell migration in Case I led to a faster expansion of the tumor. The growth of the invisible metastasis was the fastest in Case III due to presence of fibroblasts and the higher cell migration rate. The growth of the invisible metastasis in Case IV was slower than in Case III due to the lack of migration. However, the TTP did not follow this ordering, as the TTP depends on the total number of cells in all of the metastatic sites.

**Figure 16.**
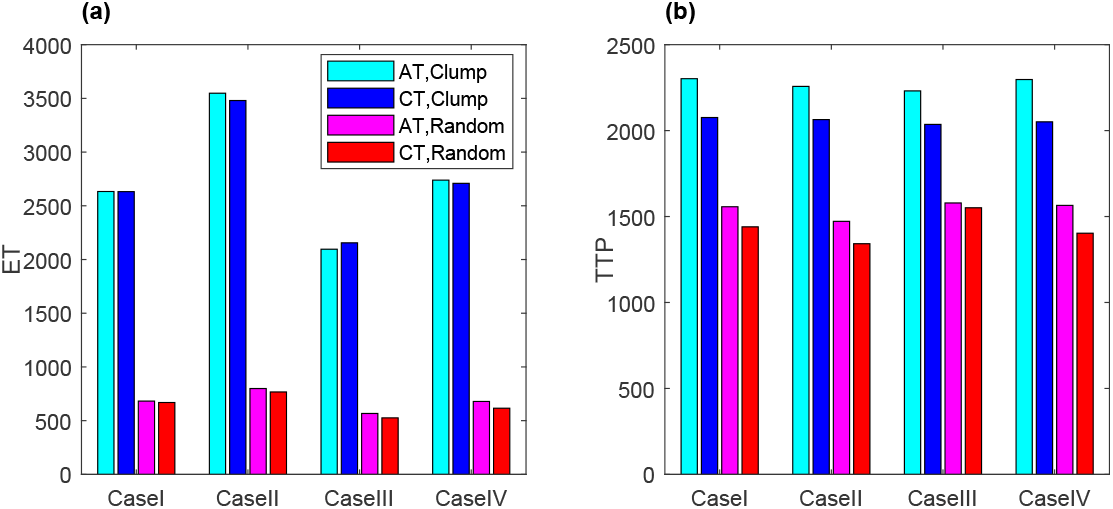
Bar chart of the ET and TTP under CT and AT for Cases I to IV with (a) clumped or (b) random invisible metastases.

## 4. Discussion

Adaptive therapy has been shown to offer delayed progression with a lower cumulative dose rate by exploiting competition between tumor cells [16]. Within tumorous tissues and throughout normal tissues, cells compete for space and survival with their neighbors. As recent studies have demonstrated, the spatial structure can shape a tumor’s evolution [19,27,28,38]. This spatial competitive aspect has been further experimentally investigated [27,39], but more work needs to be done to better understand how pre-existing tumor resistance emerges and is maintained in different spatial structures of tumors and under different treatment strategies. Different initial distributions of resistant cell populations can cause different outcomes. Depending on the locations of fibroblasts, some cancer cells can survive under therapy. To examine how the effects of the spatial structures are governed by these factors, we developed a 2D agent-based model in which the sensitive cells were randomly distributed over the domain and the resistant cells were clumped near the center of the domain, randomly distributed over the domain, or uniformly distributed over the domain. Our simulations showed that a clumped distribution of resistant cells forces high intra-species competition (R–R), leading to delayed tumor progression under therapy. The combination of high R–R competition and sustained R–S competition under adaptive therapy leads to an even longer time gain under adaptive therapy compared to continuous therapy. A reduction in R–R competition through an increase in the local carrying capacity and cell migration promotes a faster relapse.

Our analysis of the effects of the distribution of fibroblasts on resistance suggested that there may be an optimal proximity to fibroblasts for maximal tumor cell growth advantage. For resistant cells that are already competing (overlapping R-cells and fibroblasts), the fibroblast-mediated advantages of tumor progression are not significant. On the other hand, if fibroblasts are close—but not too close—to resistant cells (e.g., when Fsq encapsulates resistant cells), resistant cells on the leading edge that experience less competition can exploit fibroblast-mediated growth, leading to much faster tumor progression in both continuous and adaptive therapy. In our simulations, fibroblasts promoted sensitive cell proliferation, which unexpectedly increased the chance of drug-induced cell death because only proliferating sensitive cells can engage in cell death. During the “off” treatment in the adaptive therapy cycles, both cell types gained the same promotion promotion of by fibroblasts. Thus, the competition between the resistant cells and sensitive cells was unexpectedly reduced, resulting in a negligible benefit of adaptive therapy compared to continuous therapy.

The differential characteristics of metastatic lesions drive the evolution of tumors and the success of treatments [40–43]. A new metastatic lesion can be detected in spite of the administration of therapy. Our simulation on a virtual patient with four metastatic lesions—with one being initially undetected—predicted complex interactions between the tumor cells and fibroblasts within each metastatic lesion. Surprisingly, we demonstrated that invisible metastatic lesions can cause a rapid failure of treatments, highlighting the importance of tracking metastatic lesions during therapy. The release of a serological marker for monitoring advanced tumors, such as LDH (lactate dehydrogenase for melanoma) [44] or PSA (prostate-specific antigen for prostate cancer) [45], may be different between primary and metastatic sites or between metastatic sites [46]. Novel imaging technologies need to be developed in order to allow for frequent non-invasive monitoring of tumor burdens. Such new technologies could offer the opportunity to better understand tumors’ spatial structures.

The model presented here is an abstract representation of what might be happening in actual tumors; it focuses on spatial variations, but not how the variations arise. For example, we did not consider different microenvironmental factors, such as oxygen levels, or growth factors. The model rests on the assumption that two key tumor cell populations—drug-sensitive and drug-resistant cell populations—compete. We also assumed a uniform drug distribution, but in reality, the diffusion of a drug through a tumor tissue could result in a spatially heterogeneous drug response [7]. The adaptive strategy for the therapy used in this study considers the initial tumor volume and one threshold for stopping treatment in order to determine the on–off cycles of the treatment. However, in several studies, the maintenance and reduction of the critical volume (not necessarily the initial volume) at different levels have been reported to be beneficial [20,21,25]. We chose our modeling approach as a starting point in order to better understand how the spatial distributions of resistant cells and fibroblasts impact the outcomes of adaptive therapy.

In future studies, a few other dimensions, such as sequential dosing, alternating dosing, or fibroblast inhibitors, could be incorporated into adaptive treatment strategies [47]. Multidrug therapy was recently found to be promising by West and colleagues [23,24], but they did not consider the spatial aspects of tumors. Our simulations demonstrated that fibroblasts can cause a faster failure of adaptive therapy. In tumors, fibroblasts influence the growth of the tumor cells in a spectrum of ways [48–51]. For example, in breast cancer, fibroblasts increase the growth by secreting epidermal growth factor (EGF); furthermore, the transforming growth factor-*β* (TGF-*β*) produced by the tumor cells converts fibroblasts into myofibriblasts, which increase the secretion of EGF and thus cause even more rapid tumor progression [52]. In colon cancer, TGF-*β*1 was found to promote tumor growth by helping fibroblasts to influence tumor cells [53]. Therapies designed to target fibroblasts have been proven to be successful in cases such as liver cancer [54] and prostate cancer [55]. An adaptive therapy that combines these drugs may prolong survival with lower cumulative dose rates.

## Author Contributions

Conceptualization, M.M. and E.K.; methodology, M.M.; ; formal analysis, M.M.; investigation, M.M, J.K, C.P, E.K.; resources, J.K, C.P., E.K.; data curation, M.M.; writing—original draft preparation, M.M and E.K.; writing—review and editing, J.K., C.P.; visualization, M.M.; supervision, E.K.; funding acquisition, C.P., E.K. All authors have read and agreed to the published version of the manuscript.

## Funding

This research was funded by the National Research Foundation Korea, NRF-2019R1A2C1090219, and the KIST institutional program through grants 2Z06482 and 2Z06483.

## Conflicts of Interest

The authors declare no conflict of interest.

## Abbreviations

The following abbreviations are used in this manuscript:

CT: Continuous therapy
AT: Adaptive therapy
VNHD: Von Neumann neighborhood
TTP: Time to tumor progression
TR50: Time for resistant cells to grow to 50% of the initial tumor volume
ET: Emergence time
TG: Time gain
PPr: Progression probability

### Appendix A

#### Appendix A.1

**Figure A1.**
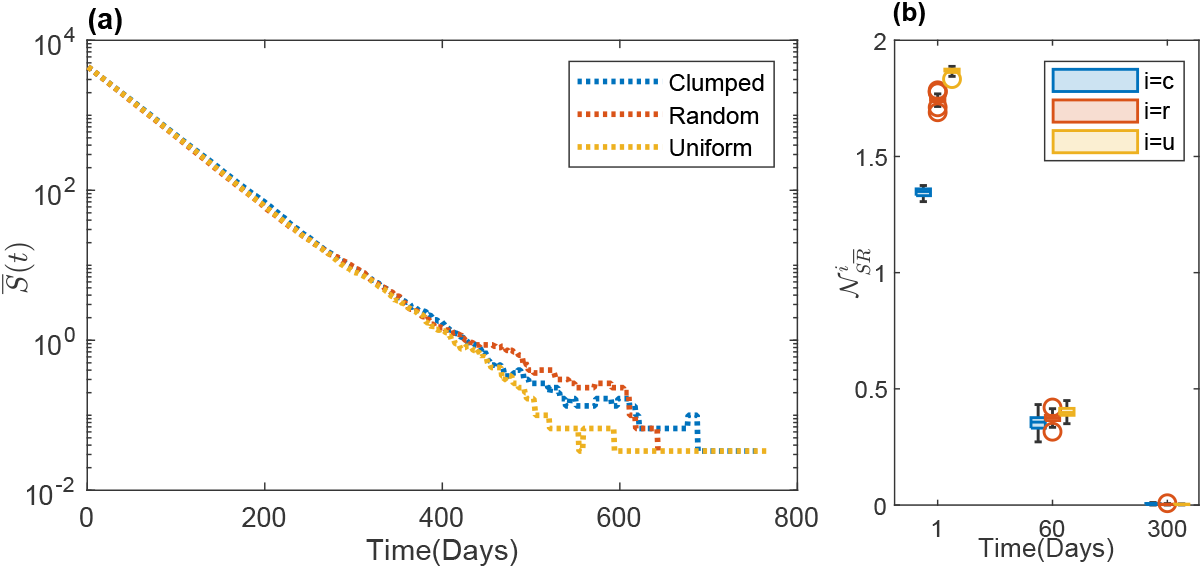
(**a**) The temporal evolution of the mean number of S-cell 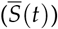 populations under continuous therapy with initial clumped, random, and uniform cell configurations is shown in a log plot, which shows very similar growth patterns among the different cases. (**b**)The average numbers of S-cells in the VNHD of an R-cell in the 30 realizations are shown as boxplots.

**Figure A2.**
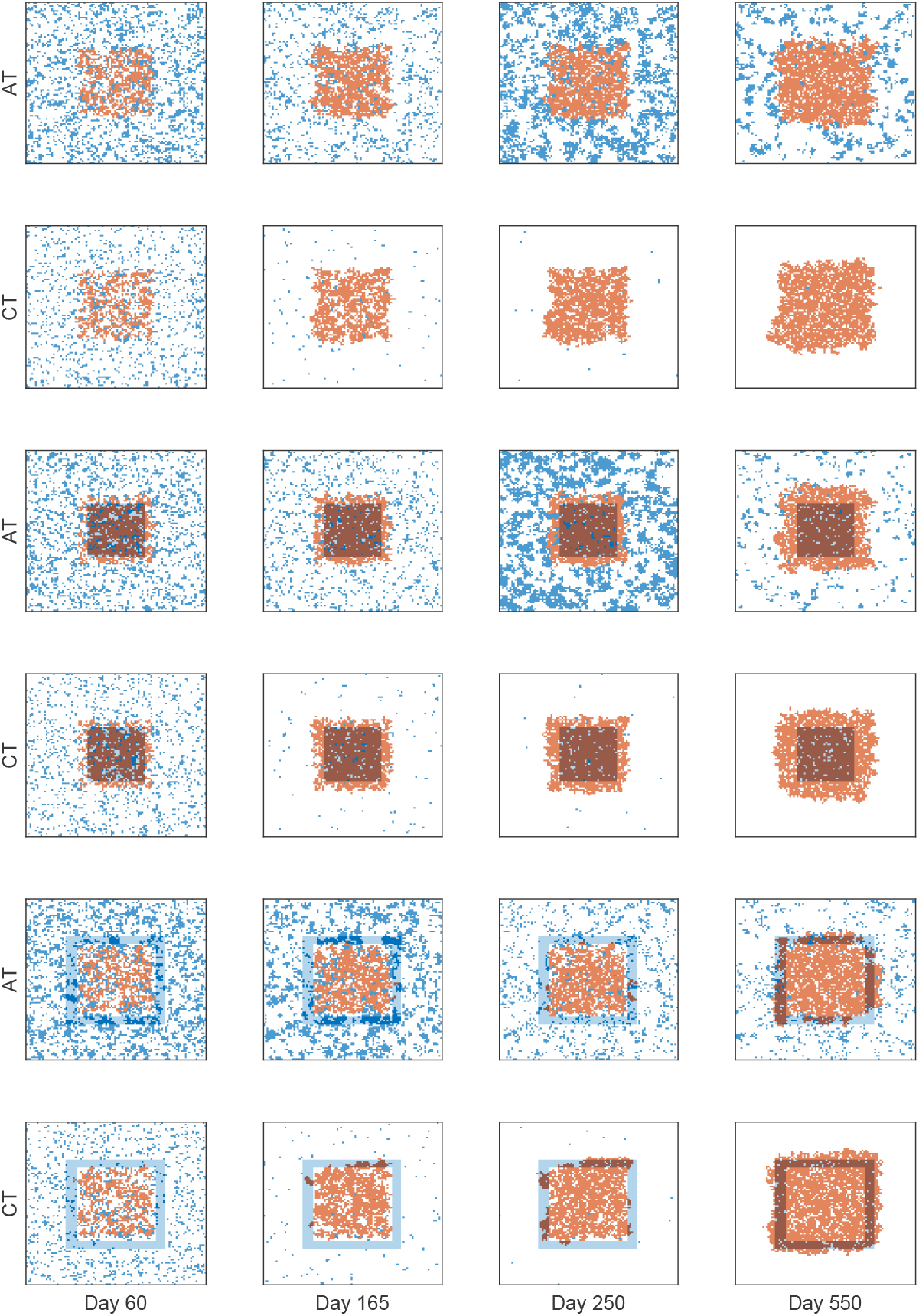
Cell configurations for fibroblast-mediated growth. The above figures show the cell configurations at different times under both AT (odd rows) and CT (even rows) for the fibroblast configurations of NoF (rows 1 and 2), FC (rows 3 and 4), and FSq (rows 5 and 6). In row 4, we see that most of the R-cells belong to the fibroblast region and, hence, experience a fibroblast-mediated increase in growth rate until the local carrying capacity is reached. In row 6, initially, none of the R-cells belong to the fibroblast region; hence, they grow at a normal growth rate. However, with time, the S-cells become sparse due to the administration of drugs, and the clump grows to reach the fibroblast region (days 165 and 250). During this time, the outer cells grow at faster rate due to the fibroblast-mediated advantages and the lower competition. When the outer cells expand past the fibroblast region (day 550) and the fibroblast region reaches its carrying capacity, the fibroblast-mediated advantages do not have an impact because of the local carrying capacity. Similarly, under AT, the R-cells mostly take advantage of the fibroblast-mediated growth in the FC configuration. However, in the FSq configuration, only a few S-cells obtain similar advantages, but no R-cells do. When the clump grows enough for the outer cells to reach the fibroblast region, the R-cell population experiences fibroblast-mediated growth.

**Figure A3.**
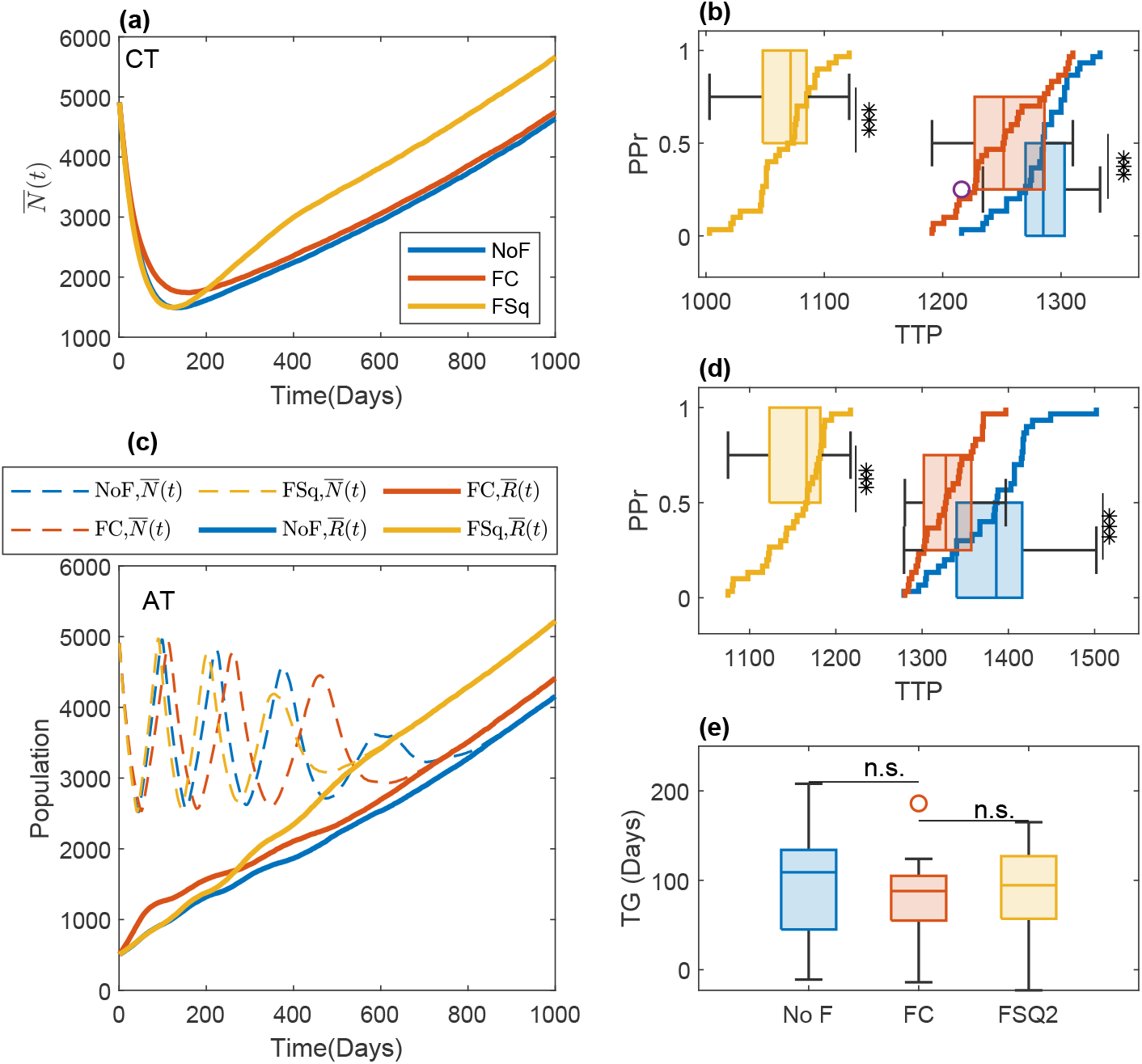
Consequences of fibroblast-mediated growth for the time to progrssion and the time gain of adaptive therapy compared to continuous therapy with a cell migration rate of *m* = 50% of the cell growth rate. (**a**) The time evolution of the mean of the total cell population 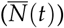 under CT in the 30 simulations is shown for the three types of fibroblast configurations—NoF, FC, and FSq—with blue, red, and yellow lines, respectively. (**b**) Boxplot of the time gain under CT in the 30 realizations, along with the progression probability (PPr) for the three types of fibroblast configurations—NoF (blue), FC (red), and FSq (yellow). (**c**) The time evolution of the average numbers of the total cell population 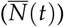, R-cells 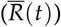, and S-cell 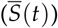 under AT in the 30 simulations are shown with dashed, solid, and dotted lines for the three types of fibroblast configurations—NoF, FC, and FSq—with blue, red, and yellow lines, respectively. (**d**) Boxplot of the time gain under AT in the 30 realizations, along with the progression probability (PPr) for the three types of fibroblast configurations—NoF (blue), FC (red), and FSq (yellow). (**e**) The blue, red, and yellow boxplots show the time gain for the three types of fibroblast configurations—NoF, FC, and, respectively. Though in the FSq configuration, a reduction in TG was observed under both CT (in (b)) and AT (in (d)), no significant differences were observed in the TG.

**Figure A4.**
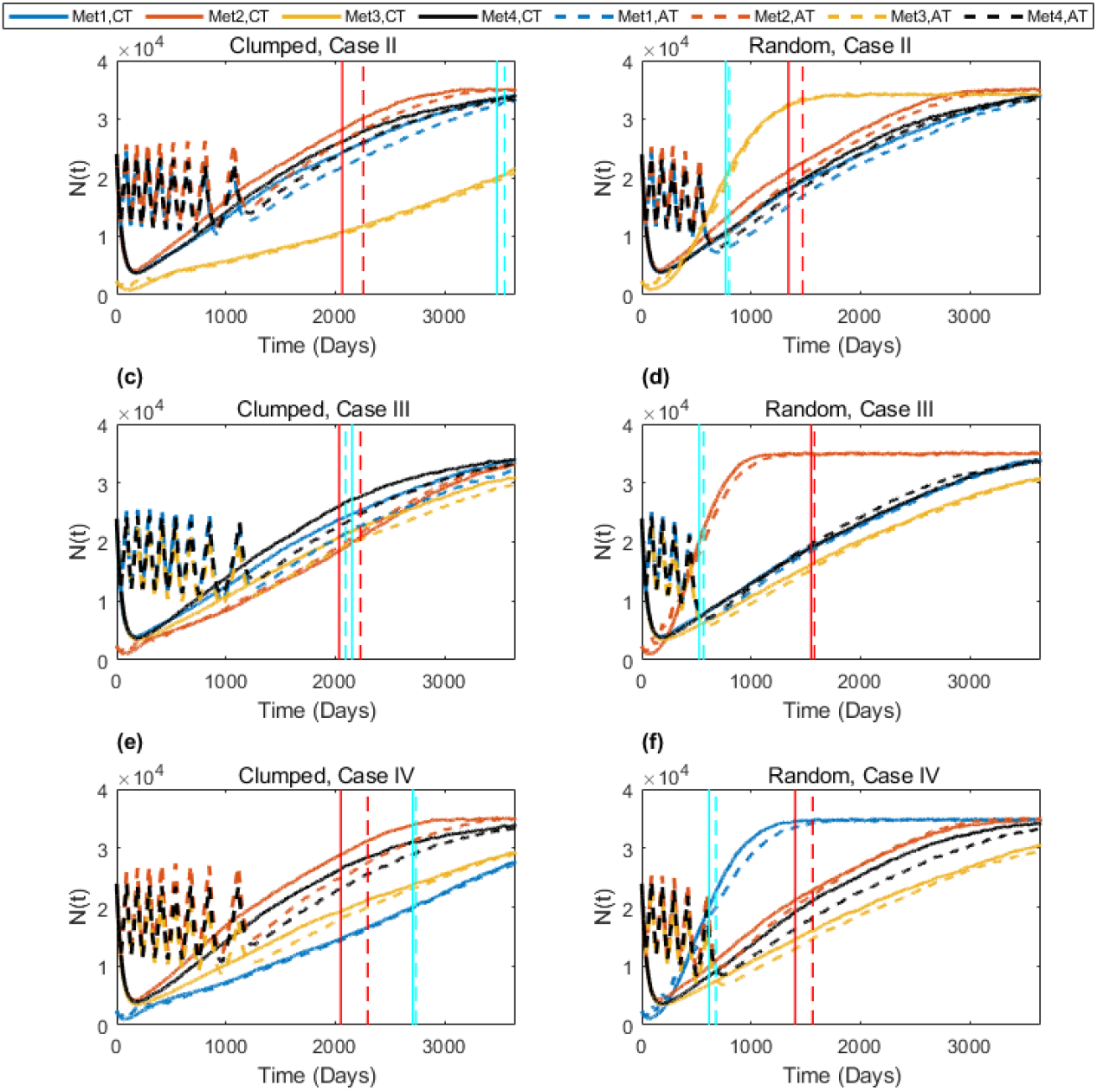
Complex dynamics of multiple metastases under AT and CT. The time evolution of the total cell population in the four metastases is shown in the sub-figures. The first, second, and third rows show the results for Cases II, III, and IV, respectively. The first and second columns show the results for clumped and random initial cell configurations in the invisible metastasis, respectively. In each sub-figure, the blue, red, yellow, and black colors show the total cell populations in metastasis 1, metastasis 2, metastasis 3, and metastasis 4, respectively; the vertical cyan lines show the emergence time (ET) of the invisible metastasis, and the red line shows the TTP. The solid and dashed lines show the results under CT and AT, respectively.

## Notes

### Competing Interest Statement

The authors have declared no competing interest.

